# TrackSig: reconstructing evolutionary trajectories of mutations in cancer

**DOI:** 10.1101/260471

**Authors:** Yulia Rubanova, Ruian Shi, Caitlin F Harrigan, Roujia Li, Jeff Wintersinger, Nil Sahin, Amit Deshwar, Quaid Morris, PCAWG Evolution and Heterogeneity Working Group, PCAWG network

**Affiliations:** Department of Computer Science, University of Toronto, Toronto, Canada; Donnelly Centre for Cellular and Biomolecular Research, University of Toronto, Toronto, Canada; Department of Electrical and Computer Engineering, University of Toronto, Toronto, Canada; Department of Molecular Genetics, University of Toronto, Toronto, Canada; Vector Institute, Toronto, Canada; Ontario Institute for Cancer Research, Toronto, Canada

**Author notes:** Various affiliations (PCAWG Evolution and Heterogeneity Working Group, PCAWG network). On behalf of the PCAWG Evolution and Heterogeneity Working Group and the ICGC/TCGA Pan-Cancer Analysis of Whole Genomes Network.

## Abstract

We present a new method, TrackSig, to estimate the evolutionary trajectories of signatures of different somatic mutational processes from DNA sequencing data from a single, bulk tumour sample. TrackSig uses probability distributions over mutation types, called mutational signatures, to represent different mutational processes and detects the changes in the signature activity using an optimal segmentation algorithm that groups somatic mutations based on their estimated cancer cellular fraction (CCF) and their mutation type (e.g. CAG->CTG). We use two different simulation frameworks to assess both TrackSig’s reconstruction accuracy and its robustness to violations of its assumptions, as well as to compare it to a baseline approach. We find 2-4% median error in reconstructing the signature activities on simulations with varying difficulty with one to three subclones at an average depth of 30x. The size and the direction of the activity change is consistent in 83% and 95% of cases respectively. There were an average of 0.02 missed and 0.12 false positive subclones per sample. In our simulations, grouping mutations by mutation type (TrackSig), rather than by clustering CCF (baseline strategy), performs better at estimating signature activities and at identifying subclonal populations in the complex scenarios like branching, CNA gain, violation of infinite site assumption, and the inclusion of neutrally evolving mutations. TrackSig is open source software, freely available at https://github.com/morrislab/TrackSig.

## 1 Introduction

Somatic mutations accumulate throughout our lifetime, arising from external sources or from processes intrinsic to the cell^1, 2^. Some sources generate characteristic patterns of mutations. For example, smoking is associated with G to T mutations; UV radiation is associated with C to T mutations^3–5^. Some processes provide a constant source of mutations^6^ while others are sporadic^7^.

One can estimate the contribution of different mutation processes to the collection of somatic mutations present in a sample through mutational signature analysis. In this type of analysis, single nucleotide variants (SNVs) are classified into 96 types based on the type of substitution and tri-nucleotide context (e.g., ACG to a ATG)^2^. Mutational signatures across the 96 types were derived by non-negative matrix factorization in the previous work by *Alexandrov et al*.^2^. Many of the signatures are associated with known mutational processes including smoking^7^, non-homologous double strand break repair^2^, and ionizing radiation^8^. The activities of some signatures are correlated with patient age^6^ and suggest their use as a molecular clock^9^. Thus, signature analysis can identify the DNA damage repair pathways that are absent in the cancer, can predict prognosis^10^ or guide treatment choice^11^.

Formally, a *mutational signature* is a probability distribution over a categorical variable representing a mutation type, where each element is a probability of generating a mutation from the corresponding type^12^. Each signature is assigned an *activity* (also called *exposure*) which represents the proportion of mutations that the signature generates. Activities for pre-defined signatures can be computed from the total mutational spectrum of a sample by using constrained regression^13, 14^.

Mutational sources can change over time^15–19^. Mutations caused by carcinogen activity stop accumulating when the activity ends^7^. Mutations associated with defective DNA damage repair, such as BRCA1 loss^1, 2^ will begin to accumulate after that loss. Recent analyses of sequencing data from single, bulk samples have reported modest changes in signature activities between clonal and subclonal populations^9, 20^ based on groups of mutations identified by clustering their variant allele frequencies (VAFs). However, the accuracy of these methods relies heavily on the sensitivity and precision of this clustering, which is typically low^21, 22^ except, in some cases, in multi-region sequencing studies^15–19, 23^.

Here we introduce TrackSig, a new method to reconstruct signature activities across time without VAF clustering. We use VAF to approximately order mutations based on their prevalence within the cancer cell population and then track changes in signature activity that are consistent with this ordering.

We use realistic simulations and bootstrap analysis to help assess the accuracy of signature activity reconstructions under a variety of different evolutionary scenarios. Using TrackSig, we have previously demonstrated^24^ that signature activities change often during the lifetime of a cancer. Here we show that these changes can be often be a more sensitive indicator of new subclonal lineages than VAF clustering.

## 2 Methods

TrackSig is designed to be applied to VAF frequency data from a single, heterogeneous tumour sample. The method consists of two stages. First, we sort single nucleotide variants (SNVs) by their estimated cancer cell fraction (CCF) that we estimate using their variant allele frequencies (VAFs) and a copy number aberration (CNA) reconstruction of the samples. Next, we infer a trajectory of the mutational signature activities over the estimated ordering of the SNVs. We estimated activity trajectory for each signature as a piece-wise constant function of the SNV ordering with a small number of changepoints. These stages are described in detail below. Note that TrackSig does not rely on any methods for clustering mutations, such as phylogeny reconstruction.

### 2.1 Ordering the SNVs

No single evolutionary model can yet explain all of the observed VAF distributions in bulk tumor samples^21, 23, 25–27^. Using a CCF-based ordering of SNVs allows TrackSig to track changes in signature activity under a variety of such models. The clonal evolution model of cancer^28^ posits that SNVs belong to one of a handful of subclones whose associated mutations all have the same CCF. Under this model, mutations from different subclones would occupy distinct regions of the CCF space, so if signature activities differ between subclones, the segments detected by TrackSig in CCF space would mark the presence of distinct subclones. Current neutral evolution models^21, 25^ assume SNVs to be unique and persistent (i.e. the *infinite sites assumption*) and SNVs with higher CCFs generally occur earlier in the tumor’s evolution. Therefore, when SNVs are sorted in order of decreasing CCFs, TrackSig’s trajectories track changes in signature activity over time.

In the following sections, we describe how the SNV VAFs are used to create a *timeline* to which TrackSig is applied. For ease of presentation, we will assume that the time of SNV occurrence increases approximately monotonically with position on the timeline. This interpretation is valid under the infinite sites assumption and either a neutral evolution model or a clonal one when all subclones are from the same branch, as is often the case in single samples^29^. In section 3.2.2, we test TrackSig’s reconstruction accuracy on simulated data that is consistent with different evolutionary models and violations to these assumptions; and in section 4.2, we discuss how to interpret TrackSig’s reconstructions when the timeline is not a faithful representation of time of SNV acquisition.

#### 2.1.1 Estimating cancer cell fraction

Estimating a SNV’s CCF requires both an estimate of its VAF and an estimate of the average number of mutant and reference alleles per cell at the locus where the SNV occurs. In TrackSig, we derive this estimate from a CNA reconstruction provided with the VAF inputs.

To account for uncertainty in a SNV’s VAF due to the finite sampling, we model the posterior distribution over its VAF using a Beta distribution:

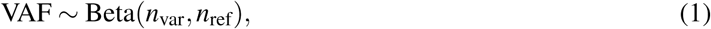

where *n*_var_ is the number of reads carrying a variant, and *n*_ref_ is the number of reference reads. To simplify the algorithm, and the subsequent sorting step, we sample an estimate of VAF_*i*_ (VAF of SNV *i*) from this distribution and use that sample as a surrogate for the distribution in subsequent calculations. An advantage of this approach is that it gives us a single ordering. With a large number of SNVs, we expect little variability in the estimated activity trajectory due to uncertainty in the VAFs of individual SNVs. With a smaller number of SNVs, multiple orderings can be sampled and the trajectories combined.

If no CNA reconstruction is available, TrackSig assumes that each SNV is in a region of normal copy number and TrackSig estimates CCFs in autosomal regions by setting:

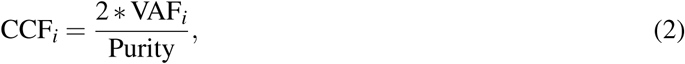

where Purity is the purity (i.e. proportion of cancerous cells) of the sample. If purity is not provided, TrackSig assumes Purity = 1.

If a CNA reconstruction is available, TrackSig uses it when converting from VAF to CCF. TrackSig assumes there is a maximum of one copy of the variant allele per cell, and thus estimates CCF by setting:

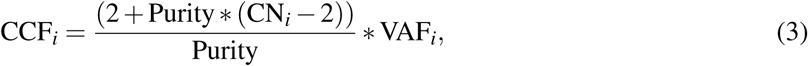

where Purity is the purity of the sample, and CN_*i*_ is the clonal copy number of the locus. If the clonal CNA increases the number of variant alleles per cell, this will lead to CCFs larger than one. As such, these cases are easily detected and corrected. Specifically, if the observed VAF is greater than 50% due to finite sampling noise, or there is more than variant allele per cell, the CCF_*i*_ calculated above could be larger than 100% which cannot be correct. As such, in most cases, we will set CCF_*i*_ to be the minimum of equation 3 and 100%. When we do not do so, we will sometimes refer to these estimates as *(estimated) average number of mutant alleles per cell* to avoid confusion.

In regions of subclonal CNAs, estimating CCF requires a phylogenetic reconstruction in order to determine whether the subclonal CNA influences the number of variant alleles in the affected cells^22, 30^. As such, when computing SNV ordering, by default, TrackSig filters SNVs in these regions out in order to avoid this time consuming operation. However, TrackSig can include these SNVs if provided with CCF estimates for them from methods that do consider subclonal CNAs^22, 30, 31^.

TrackSig sorts SNVs in order of decreasing estimated CCF and uses the rank of the SNV in this list as a “pseudo-time” estimate of its time of appearance. Note that this estimate will have a non-linear relationship to real time, if the overall mutation rate can vary during the tumour’s development. If some of the SNVs can be interpreted as clock mutations, an SNV’s rank can be converted into an estimate of real time^9^.

#### 2.1.2 Constructing a timeline

To derive an estimate of the activity trajectory, TrackSig converts the SNV ordering into a set of time points with non-overlapping subsets of the SNVs. We do this for two reasons. First, stable estimation of signature activities requires a minimum number of mutations. By binning mutations into time points and requiring a minimum number of time points per segment, TrackSig enforces a minimum of 100 mutations per segment. Also, the time complexity of TrackSig scales with the number of time points. So by binning mutations, we can speed up TrackSig. By default, we set the bin size to 100 but the user can change this setting to as low as 1. As we show in section 3.2.2, TrackSig’s signature activity reconstructions are relatively insensitive to the choice of bin size.

TrackSig first partitions the ordered mutations into bins and interprets each bin as one time point. The *timeline* of the cancer is the collection of the time points. TrackSig reports signature activity trajectories as a function of points in the timeline. We emphasize that TrackSig does not use any information about subclones when partitioning the SNVs and that TrackSig only uses CCFs for the SNV from a single sample.

### 2.2 Computing activities of mutational signatures

To estimate activity trajectories, TrackSig partitions the timelines into segments containing one or more time points. Within each of these segments, it estimates signature activities using mixture of discrete distributions. Full details of the model are provided in the appendix A. In brief, TrackSig models each signature as a discrete distribution over the *K* types and it treats the mutation count vector over the *K* types as a set of independently and identically distributed samples from a mixture of the discrete distributions corresponding to each signature. By default, TrackSig uses single base trinucleotide signatures^2^ and *K* = 96, however TrackSig can use any mutation type labelling scheme, so long as it is given appropriate signatures as input. The mixing coefficients of these distributions are interpreted as their activities for the mixture model that produced the set of mutations. TrackSig fits these activities using the Expectation-Maximization algorithm (EM)^32^, as done by other signature activity estimation methods^33^.

### 2.3 Detecting changepoints

TrackSig identifies changepoints in the timeline where there are discernible differences in the activity of mutations in the time points before and after the changepoints. Specifically, the changepoints partition the timeline into segments of mutations with approximately constant activities. TrackSig fits activities for this set using EM algorithm as described above. This procedure generates piece-wise constant activity trajectories for each signature. To select changepoints, we adapt Pruned Exact Linear Time (PELT)^34^, an optimal segmentation algorithm based on dynamic programming. We impose a complexity penalty at each time point that is equivalent to optimizing the Bayesian Information Criteria (BIC) (see Supplement B for details). To reduce variance in our estimates of the signature activities, we do not allow partitions to be smaller than 100 mutations.

We compute the BIC criteria the following way. Changepoints split the timeline into (# changepoints + 1) segments. In each segment, TrackSig fits the signature activities, which have to sum to one. Therefore there are (# signatures −1) free parameters per segment, or (# changepoints + 1)·(# signatures −1) free parameters in total. As such, BIC objective takes the following form:

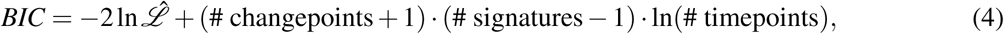

where 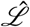 is the likelihood of the current model.

### 2.4 Correcting the timeline and segment count

#### Cancer cell fraction greater than 1

If the number of variant alleles per cell is increased by a clonal copy number change, TrackSig’s CCF estimates might be greater than 1. To correct for this, when displaying activity trajectories, it merges all the time points that have average CCF ≥ 1 into one time point. As such, the first time point can contain more than 100 mutations. To determine a signature activity at this new time point, TrackSig simply takes an average activity of all merged time points (those having CCF ≥ 1).

#### Number of distinct subclones

To compute the number of distinct subclones, we adjust the number of detected changepoints to correct for overlap in the CCF space of mutations from different subclones. Consider the case of two subclones whose mutations overlap substantially in CCF space. In this case, TrackSig might find three segments instead of two: one with signatures activities reflecting the first subclone; another with activities reflecting a mixture of the two subclones; and last with activities reflecting the second subclone. If this happens, then the direction of change of all signatures will be the same in the two changepoints. As such, when counting the number of distinct subclones, we treat each such pair of changepoints as one subclone boundary. Such a situation only occurs in 2.6% of 2,552 PCAWG tumour samples to which we applied TrackSig; in 77% of those cases we remove a single changepoint.

### 2.5 Bootstrapping to estimating activity uncertainty

TrackSig estimates uncertainty in the activity estimates by bootstrapping the mutations and refitting the activity trajectories. Specifically, it takes the random subset of *N* mutations by sampling uniformly with replacement from the *N* unfiltered SNVs in the sample under consideration. Using the pre-assigned CCF estimates, we sort the SNVs in decreasing order, as above, re-partition them into time points and recompute activity estimates. The trajectories obtained from bootstrapped mutation sets have the same number of time points, however the average CCF for each time point can change. We use these bootstrapped trajectories to compute uncertainty estimates for the sizes of activity changes.

## 3 Results

In this paper we perform the realistic simulations to evaluate TrackSig’s performance at reconstructing signature activities, detecting the number of mutation clusters and correctly placing the changepoints under different scenarios including violations of TrackSig’s assumptions.

TrackSig was applied to the 2,552 whole-genome sequencing samples with more than 600 SNVs contained within the white and grey lists of the Pan-cancer Analysis of Whole Genomes (PCAWG) group. Here we provide methodological details of TrackSig’s use on real data (PCAWG). The analysis of signature trends and relation of changepoints found by TrackSig to subclonal boundaries is described elsewhere^24^.

### 3.1 Choice of mutation signatures

By default, following *Alexandrov et al*.^2^, we classify mutations into 96 types based on their three-nucleotide context. Point mutations fall into 6 different mutation types (i.e., C -> [AGT] and T -> [ACG]) excluding complementary pairs. There are 16 (4*4) possible combinations of the 5’ and 3’ nucleotides. Thus, SNVs are separated into 96 (*K* = 16*6 = 96) types. However, TrackSig can use different mutation type labelling schemes, so long as the signatures and the mutation types are provided as input.

Within the context of PCAWG, we use the set of 48 single-base signatures (SBS) developed by PCAWG-Signature group. The first 30 of those signatures are slightly modified versions of original signatures defined by *Alexandrov et al*.^2,12^ and have the same numbering and interpretation. The original 30 signatures are described at COSMIC^1^. Signature analysis methods, including TrackSig, fit activities for only a subset of the signatures. These signatures are called the *active* signatures. The activities for the non-active signatures are clamped to zero. For example, SBS 7 has been detected almost exclusively in skin cancers and likely describes mutations caused by UV light^2^. As such, it is only assigned active status in skin cancers. In our analysis, we use the active signatures reported by PCAWG-Signature group. For analyses based on COSMIC signatures, one can use active signatures per cancer type as provided on COSMIC website. TrackSig can also be used to automatically select active signatures, as described in a later section.

### 3.2 Simulations

We tested sensitivity of TrackSig in multiple error scenarios using simulated data with known ground truth. First, because signatures overlap in the mutation types that they can produce, we first test reconstruction accuracy when SNVs are accurately assigned to the time points to assess errors due to inability to correctly assign signature activity. In section 3.2.1 we describe these *non-parametric simulations*.

In section 3.2.2, we assess reconstructions when the mutation ordering is inferred based on mutation VAF. In this scenario, reconstruction errors can occur when i) CCF estimates are inaccurate and, ii) there are two SNV clusters which overlap in CCF space but have different signature activity profiles. In the latter case, SNVs from both clusters will be located in the same or adjacent time point bins and will have a mixture of signature activity profiles from two clusters. To test reconstruction errors in these two scenarios, we produce *clonal evolution simulations* where we sample the VAF data from a clonal evolution model with binomial sequencing noise. To simulate the VAF detection limit for mutations imposed by somatic mutation calling, we remove any mutation with fewer than three variant reads.

Finally, as part of clonal evolution simulations, we assess model misspecification error by introducing violations of the assumptions of infinite sites and the relationship between CCF and timing of mutation occurrence. Also, in some simulations, we introduce mutations under neutral selection (i.e., neutrally evolving mutations). We computed the number and VAFs of these mutations using the model and effective mutation rates derived by Williams et al^25, 26^ as detailed in Appendix C.5. As a baseline, we compare TrackSig’s reconstruction error to the widely used strategy: first assigning SNVs to clusters based on VAF, then computing mutation signatures activities from the assigned mutations within each cluster.

#### 3.2.1 Non-parametric simulations

In the non-parametric simulations, we test the ideal scenario when SNVs are correctly ordered and assigned to the time point bins. Here we want to access the ability of TrackSig to reconstruct signature activities from the distribution of mutation types and place changepoints at the correct locations.

Each simulation has 50 time points, each time point is a bin of 100 mutations. This corresponds to the average number of somatic mutations detected in PCAWG. Each sample also contains four active signatures. Two of those signatures are 1 and 5, which are nearly always active in the PCAWG samples. For the remaining two signatures, we test all 1035 possible combinations of the other 46 signatures.

We generate simulations with 0 to 3 changepoints that are placed randomly on the timeline. For each segment on the timeline, we sample signature activities from a uniform distribution over activity vectors. Finally, we sample 100 mutation types per time point from the discrete distribution derived using the sampled activities as mixing coefficients for the four signatures.

Next, we run TrackSig on the simulated data and compare the reconstructed activity trajectories to the ground truth. We remove changepoints with small change, that is, where activities of all signatures change by less than 5% in reconstructed trajectories. This threshold is derived in section 3.3.1 from permutation analysis.

We computed the absolute difference between predicted activities and the ground truth at each time point and take the median across all time points and all four signatures. We called this the median activity difference per simulation. On the simulations with no changepoints, the median of these median per simulation differences is 0.7%. On simulations with 1 to 3 changepoints, this median increases slightly to 2%. The cumulative distribution of the median per simulation differences is shown in fig. 2.

**Figure 1.**
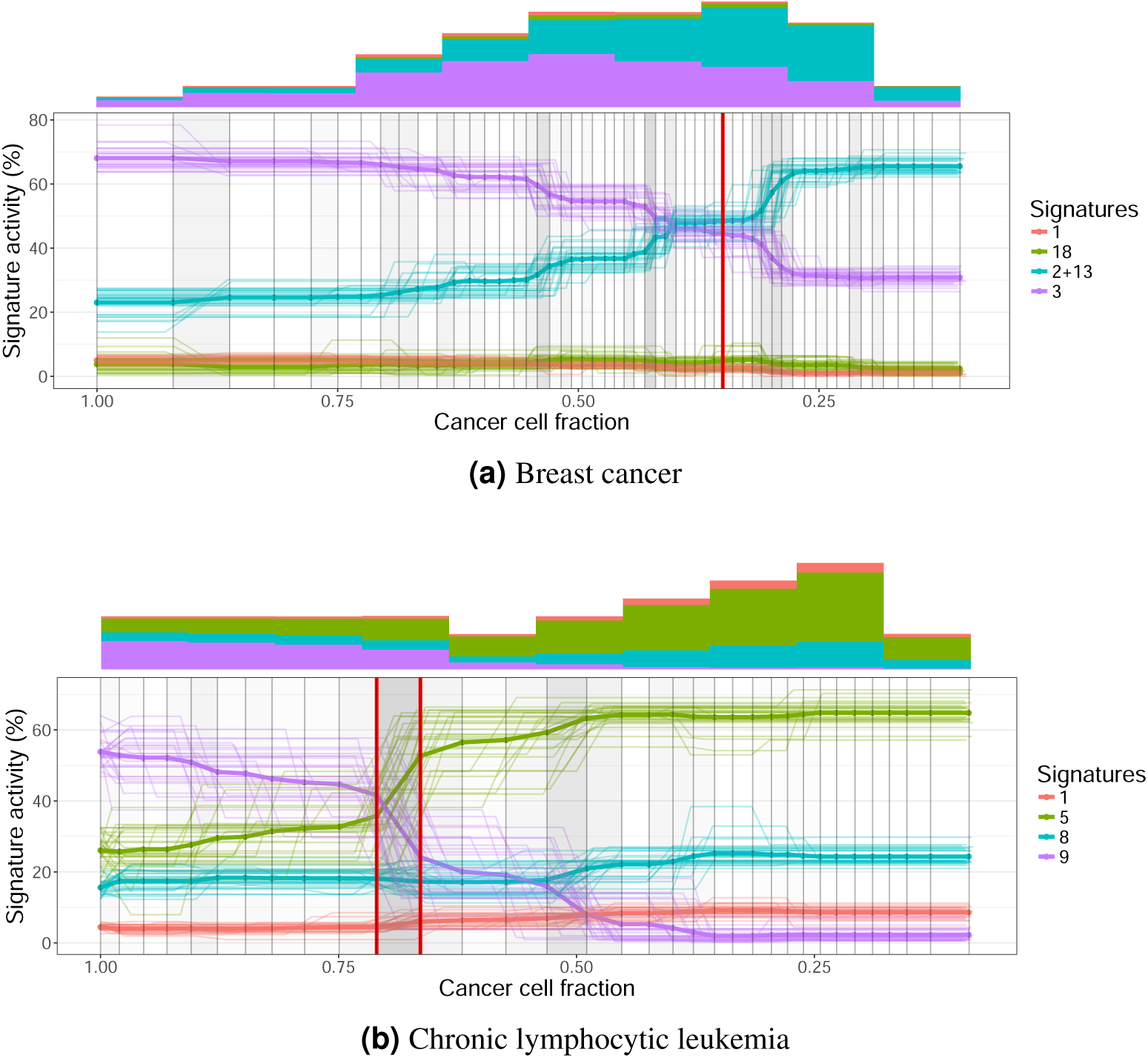
Activity trajectories for two samples. Each plot is constructed from VAF data from a single tumour sample. Each line is an activity trajectory that depicts inferred activities for a single signature (y-axis) as a function of decreasing CCF (x-axis). The thin lines are trajectories from each of 30 bootstrap runs. The bold line depicts the mean activities across bootstraps. The vertical lines indicate time points in the original dataset, and are placed at the average CCF of their 100 associated mutations. Changes in activity trajectories are not necessarily aligned with vertical bars because mean CCFs of time points change across bootstraps. Frequency of changepoints between two vertical bars is indicated by shade, the darker shades indicate higher density of changepoints. Subclonal boundaries found by PCAWG consensus clustering^24^ are shown in red vertical lines. These boundaries are not used in trajectory calculation and are only shown for comparison. Histograms show the mutation counts per signature in fixed width intervals of CCF. **(a) Breast cancer sample** In clonal signatures remain constant with dominating signature 3 (associated with BRCA1 mutations). In the subclone activity to signature 3 decreases and is replaced by SNVs associated with APOBEC/AID (signatures 2 and 13). **(b) Chronic lymphocytic leukemia sample** Signature 9 (somatic hypermutation) dominates during clonal expansion and drops from 55% activity to almost zero in the subclone. Signature 5 compensates for this change.

**Figure 2.**
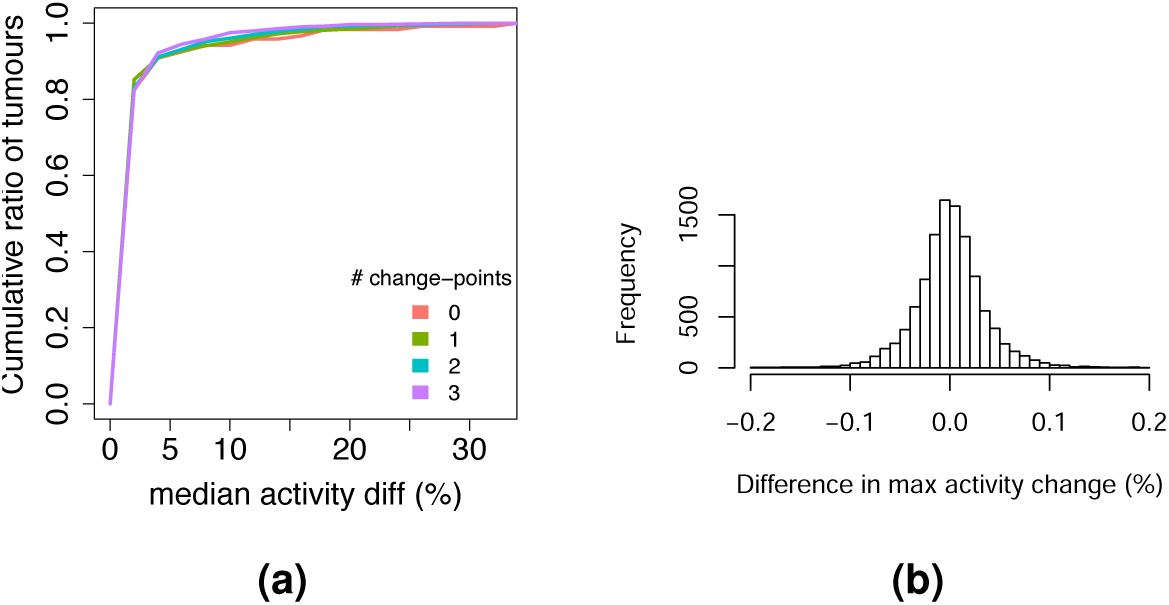
Results on non-parametric simulations. **(a)** Median activity difference between the reconstructed trajectories and the ground truth. Lines correspond to the simulations with 0, 1, 2 or 3 changepoints. The median in computed across all signatures and time points in the sample. **(b)** Distribution of Maximum Activity change (MAC) discrepancies between between estimated activities and ground truth.

For the PCAWG data, we report the maximum activity change (MAC) across activity trajectory^24^. The maximum change is the difference between maximum and minimum activity across all time points in a sample. We also report the direction of change (down if maximum occurs before minimum and up otherwise). Here, we evaluate TrackSig’s accuracy in these estimates on the simulated data. The MAC discrepancies between the estimated and ground-truth trajectories is less than 5% in 83.2% of cases across all signatures in all simulations (fig. 2b).

To compare the direction of the activity change, we divide signatures into those with: decreasing activity, increasing activity, and no activity change (i.e., max change is less than 5%). The direction of maximum change is consistent in 95.2% of all signatures across all simulations.

To compute number of false positives and false negatives, we count a true positive detection if at least one of predicted changepoints occur with three time points of an actual one. A false negative is when no predicted changepoints are within three time points of an actual change. This criteria is identical to the one we use to evaluate whether a changepoint supports a subclonal boundary^24^. We deem a predicted changepoint a false positive if it occurs more than three time points away from the closest actual changepoint.

Table 1 shows the percentage of simulations where we observe a certain number of false positives. On average, there are 0.12 false positives per simulation and 0.02 false negatives on average per simulation.

**Table 1.** Results on non-parametric simulations. False positives and false negatives versus changepoints in the ground truth. Each cell shows the percentage of simulations that have certain number of false positives/negatives (normalized within the column). See main text for definition of positive and negative time points. The last row of the table shows the average number of false positives per simulation.

|  |  | # true changepoints |  |  |  |
| --- | --- | --- | --- | --- | --- |
|  |  | 0 | 1 | 2 | 3 |
| # false positive changepoints (FP) | 0 | 0.909 | 0.896 | 0.9 | 0.889 |
|  | 1 | 0.06 | 0.083 | 0.087 | 0.106 |
|  | 2 | 0.024 | 0.019 | 0.011 | 0.005 |
|  | 3 | 0.005 | 0.002 | 0.002 | 0 |
|  | 4 | 0.002 | 0 | 0.001 | 0 |
| Avg # of FP per simulation |  | 0.130 | 0.128 | 0.118 | 0.116 |
| # false negative changepoints (FN) | 0 | 1 | 0.992 | 0.962 | 0.947 |
|  | 1 | 0 | 0.008 | 0.038 | 0.049 |
|  | 2 | 0 | 0 | 0 | 0.003 |
| Avg # of FN per simulation |  | 0.0 | 0.008 | 0.038 | 0.058 |

**Table 2.** Simulation results. Predicted changepoints versus changepoints in the ground truth. Each cell shows the percentage of simulations which have certain number of estimated changepoints (normalized within a column). Note that due to smoothing, there might be several predicted changepoints that correspond to the same changepoint in the ground truth. In this case, predicted changepoints have to be located no more than 3 time points away from the ground truth.

|  |  | # true changepoints |  |  |  |
| --- | --- | --- | --- | --- | --- |
|  |  | 0 | 1 | 2 | 3 |
| # predicted changepoints | 0 | 0.91 | 0.004 | 0 | 0 |
|  | 1 | 0.061 | 0.9 | 0.019 | 0.001 |
|  | 2 | 0.024 | 0.078 | 0.898 | 0.037 |
|  | 3 | 0.006 | 0.02 | 0.075 | 0.861 |
|  | 4 | 0.002 | 0.001 | 0.009 | 0.091 |
|  | 5 | 0 | 0.001 | 0.001 | 0.002 |

#### 3.2.2 Clonal evolution simulations

Generating realistic simulated data requires making some assumptions about how tumours evolve. In this section, we simulate VAF data consistent with clonal evolution theory^28^, with some small violations. Specifically, every mutation belongs to one of a small set of subclones. If the CCFs of each mutation could be estimated precisely, then mutation CCF data consistent with clonal evolution theory would have signature activities that are piece-wise constant functions of CCF. These VAF data are thus consistent with all previous work estimating signature activities.

We also include some simulations that violate the clonal evolution assumptions and test TrackSig’s robustness to these violations. We performed six different simulations, described briefly here (see Suppl C for details). We generated 100 simulations of each type.

##### One- and two-clusters

First, we aim to evaluate false negative and false positive rates of identifying subclones via TrackSig. We simulated VAF data from a) one clonal population and no subclones b) one clonal population and one subclone with a variety of CCF values sampled from a uniform distribution, assuming a linear clonal tree. We sample the variant allele counts for the mutations in accordance to the cluster CCFs from a binomial distribution. We create simulations with four signatures – age-related signatures 1 and 5 and two other randomly-chosen signatures, which we will refer to as A1 and A2. The activities of A1 and A2 are sampled uniformly in each clone, under the constraint that at least one of them has a signature change of at least 30%. We sample mutation types from the signature mixture treating it as a multinomial distribution. We simulated mean read depths of 10, 30 and 100.

##### One- and two-clusters, neutral evolution

We sampled mutation VAFs as per the previous paragraph but we also added some neutrally-evolving mutations to the clonal cluster. We determined the number of neutral mutations to add and sampled their VAFs according to the model from the Williams et al (2016)^25^ (see Appendix C.5). The signature activities for the neutral mutations were the same as those of the other mutations associated with the clonal cluster.

##### Branching

To assess TrackSig’s accuracy when the timeline does not reflect the ordering of acquistion of SNVs, we generated VAF data from a branched phylogeny. In branching simulation we generated VAF data assuming a branching clonal tree with two subclones. We force the sum of subclonal CCFs to be less than 1, otherwise the infinite sites assumption will be violated^1^. A later occurring subclone with a different signature activity profile has a higher CCF than a subclone with a profile matching the clonal fraction. We sample mutation VAFs and mutation types for the clusters similarly to the one- and two-cluster cases.

##### CNA gain

We next assess TrackSig’s accuracy for reconstructing activity changes when SNV VAFs are affected by a CNA. We generated VAF data with a clonal CNA gain affecting 10% of the SNV VAFs. In 5% of mutation the CNA gain is affecting the mutant allele and in 5% the CNA gain is affecting the reference allele. This simulation is created similarly to branching with three clusters. The difference is that we modify the probability of sampling a mutant allele to consider the mutant and reference copy numbers.

##### Violation of infinite site assumption

Finally, we create simulations with violation of infinite site assumption, where the same mutations independently occurred in two branched subclones. To model this, we set the CCFs of 3% of mutations to be equal to the sum of CCFs from the two subclones.

##### Comparison to SciClone + DeconstructSigs

We compared results of TrackSig to the widely used approach of first clustering mutations by CCF and then inferring signature activities within each cluster^15–19^. We perform the clustering using SciClone^35^. We do a hard assignment of mutations to clusters detected by SciClone, and use DeconstructSigs^13^ to estimate signature activities within these clusters. We report the results with two clustering methods in SciClone: Beta mixture model (BMM, default) and Beta-binomial mixture model (Binomial BMM). Note that the beta-binomial is an exact match to the noise model used in our data simulation, so we expect excellent performance from SciClone. We use the Beta model to simulate inaccuracies due to incorrect noise model specification.

##### Simulation results

We compared TrackSig and Sciclone + DeconstructSigs pipeline (hereafter SciClone for brevity) against the ground truth in the simulation across the seven simulation types and depths 10, 30, and 100. First, we computed the median activity error over all mutations in the five types of non-neutral simulations, see fig. F.7. In fig. F.8 we compared the errors in the neutral one- and two-cluster cases. Across all depths, the majority (83.6%) of TrackSig reconstructions have < 0.05 median error with ground truth versus 55.0% in SciClone. On average TrackSig’s activity error is 4.5%; SciClone’s is 6.9% across all seven simulations. Measuring activity error using KL divergence gives similar results (fig. F.4), and TrackSig’s accuracy is relatively insensitive to bin size at multiple depths (see suppl. figs. F.5, F.6)

We then compared methods based on their ability to detect subclones. Fig. 3b shows the percentage of simulations when each method predicted correct number of subclones for depth 30 (see suppl. fig. F.2 for depths 10 and 100). For this comparison, we used two different noise models with SciClone, the binomial-beta model which is an exact match to how the simulated data are generated, and the beta model, which is not and which was the model used for computing the reconstruction error above.

**Figure 3.**
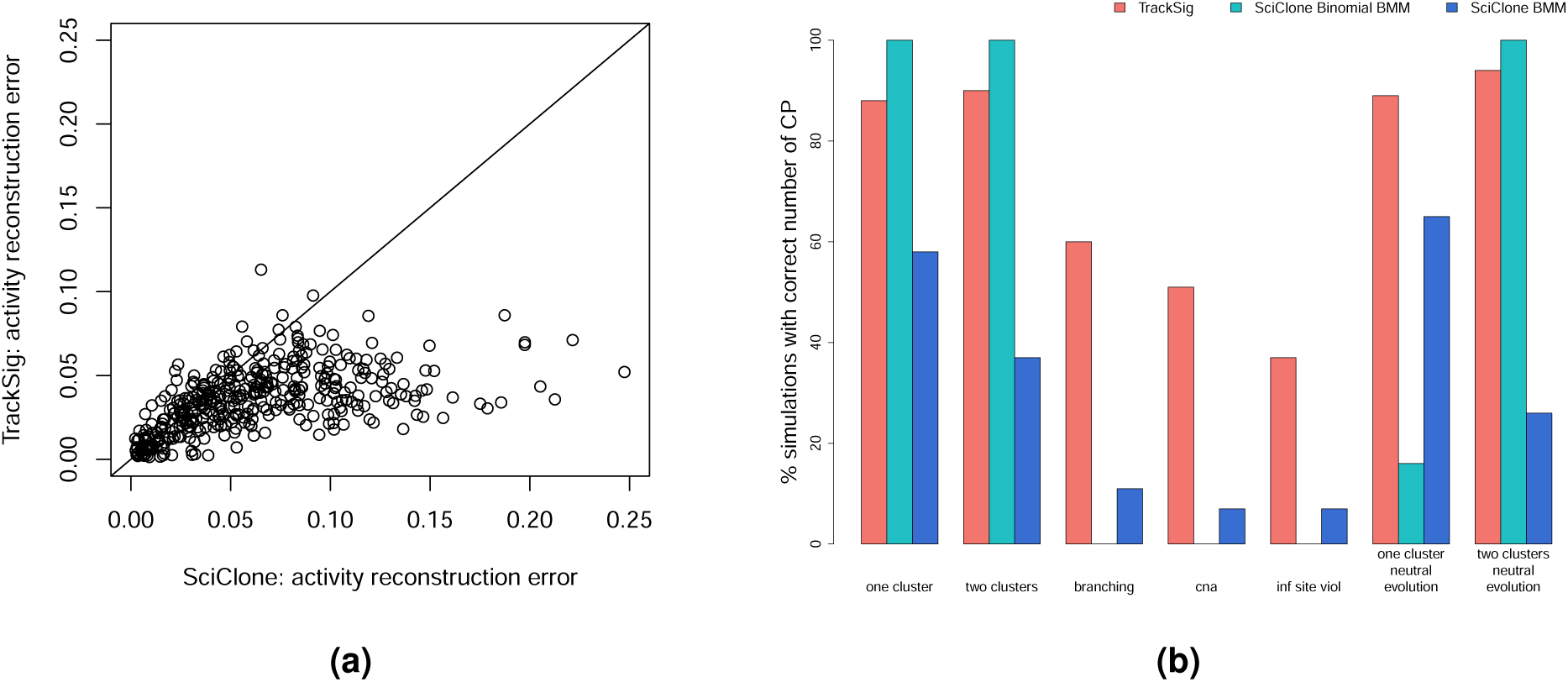
Comparison of TrackSig and SciClone baseline on clonal evolution simulations. **(a)** Scatterplot of median activity errors (i.e., absolute activity difference) on all depth 30 simulations. Mean activity error: TrackSig 3.5%, SciClone 6.2%. **(b)** Grouped barplot shows proportion of simulations where each method predicts the correct number of subclones for different simulation types as indicated on X-axis label. Different SciClone bars indicate different noise model selections. Results are for the simulations of average depth 30. Results for depths 10 and 100 are shown in figure F.2.

As expected, the SciClone binomial-beta performs nearly perfectly on the depth 30 simulations which match the assumptions of this model (Fig 3b, one-cluster, two-clusters). However, the binomial-beta model is fragile, and performs poorly when its assumptions are violated (Fig 3b, branching, cna, inf sites viol, one cluster neutral evolution). The beta model does not perform well under the ideal situation, but it is better than the binomial model when the clonal evolution assumptions are violated. In contrast, TrackSig retains the approximately 10% false positive rate in calling subclones observed in the non-parametric simulations. TrackSig’s high performance is maintained also in the neutral mutation simulations. In the other violation scenarios TrackSig has much better performance than either of the two SciClone noise models.

Figure F.2 shows that at depth 100, TrackSig has approximately 90% accuracy in all scenarios except the two cluster neutral evolution simulation. This scenario is particularly difficult because the clonal cluster is split with about 500 neutral mutations from the clonal lineage clustered at the VAF detection limit; so the mutation type distributions actually have two clear changepoints: one going from the clonal lineage to the subclonal one, and then another returning to the clonal (Fig 4a). It may be possible to detect this error in post-processing (see Section 4.2). The depth 10 simulations are particularly challenging as well, because the VAF distributions of the clonal and subclonal clusters overlap substantially, making it difficult to detect multiple clusters. Here we see that TrackSig is an even more sensitive detector of a second cluster than the correct SciClone binomial model, see Fig 4b. None of the methods do well in the branching, cna, and infinite site violation scenarios because they require the detection of three clusters. Performance in the neutral evolution scenarios here matches that in the non-neutral ones because there are very few neutral mutations above the VAF detection limit.

**Figure 4.**
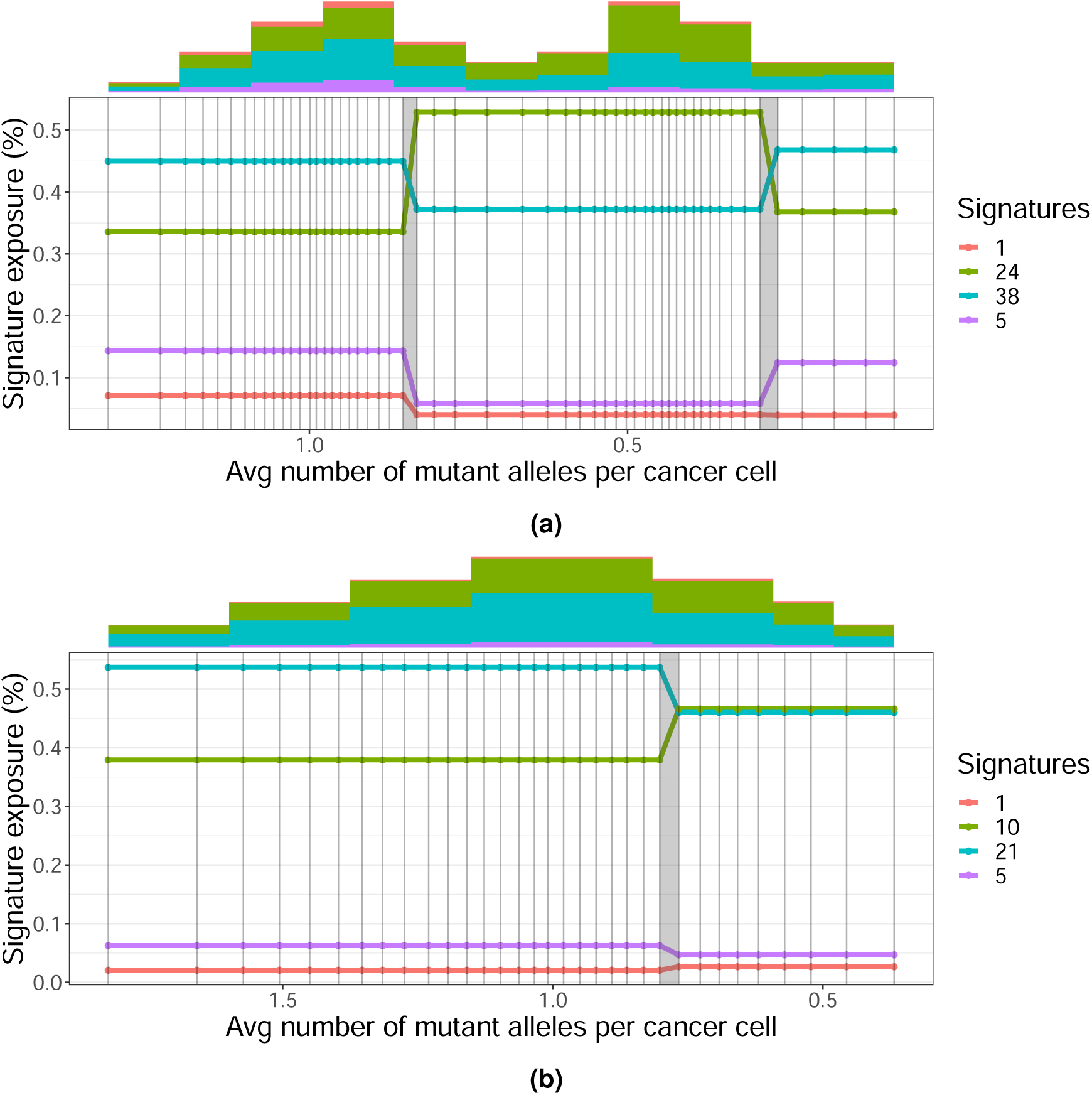
TrackSig reconstruction examples on clonal evolution simulations. **(a)** Simulated data was generated with two clusters and clonal neutral mutations at read depth 100. TrackSig incorrectly places a changepoint before a cluster of neutral mutations from the clonal lineage near the VAF detection limit. However, because the signature activities match those in the clonal cluster, this error could be detected and corrected in post-processing. **(b)** Simulated data was generated with two clusters at read depth 10. TrackSig correctly identifies one changepoint. Although the simulation contains two clusters, there is only a single mode of CCF, thus making CCF-cluster-based detection of subclones impossible. However, the histogram on top shows that there are differences in mutation type distributions between the left and right tails, permitting TrackSig to correctly identify a changepoint. Both figures use an expanded x-axis that shows the whole spread of estimated CCF, this is indicated with a change in the x-label descriptor.

In summary, TrackSig is more robust to violations of the clonal evolution assumptions that are made by most subclonal reconstruction algorithms (see^23, 26^).

### 3.3 Methodology on real data

#### 3.3.1 Choosing a significance threshold

We analyze the variation of signature activities on PCAWG data across time and across samples. We compute the maximum change of the signatures in each sample, which is simply the difference between maximum and minimum activity of the signature. To assess whether a signature change is statistically significant, we permute the mutations in each sample and run the trajectory estimation on the permuted set. Since permuted mutations are not sorted in time, we expect no change in the activity trajectories over time. The maximum activity change that we observe on permuted set of mutations does not exceed 5% in any sample. Therefore, we only consider signature changes above 5% to be significant (Fig. 5).

**Figure 5.**
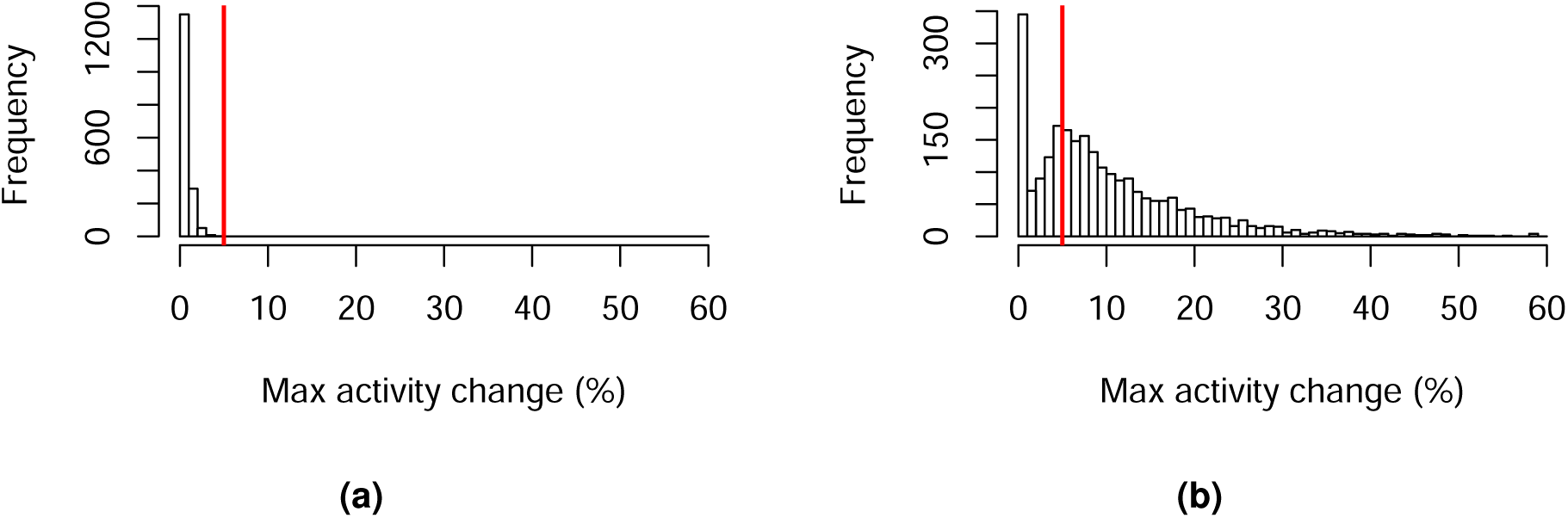
Distribution of maximum signature activity changes across 2486 PCAWG samples. The red line shows the threshold of 5%, above which we consider changes to be significant. **(a)** Changes on random orders of mutations where we don’t expect to see change in activities. **(b)** activity changes in TrackSig trajectories across all samples (on mutations sorted by CCF). Frequency axis shows the number of samples where we observe the certain activity change.

#### 3.3.2 Bootstrapping

We assess the variability in activity trajectories by performing bootstrap on the PCAWG data. We sample mutations with replacement from the original set and re-calculate their activities and changepoints. We perform 30 bootstrap runs for each sample. Fig. 1 shows examples of bootstrapped trajectories from two samples (breast cancer and leukemia).

Signature trajectories calculated on bootstrap data are stable. The mean standard deviation of activity values calculated at each time point is 2.9%. We also evaluate the consistency of signature changes across the entire activity trajectory: size of signature change and location of the changepoint. The mean standard deviation of the *signature change* is 5.3% across the bootstraps. This standard deviation does not exceed 5% in 55.8% of samples (does not exceed 10% in 94.3% of samples, fig. F.1).

In TrackSig the number of changepoints calculated during activity fitting does vary across bootstrap samples. We observe 1.02 standard deviation in the number of changepoints. To assess the variability in the location of the changepoints, we matched nearby changepoints between bootstrap samples and measured their average distance in CCF. Because the number of changepoints can change between samples, as a reference, we randomly choose one of the samples that has a number of changepoints equal to the median number of changepoints among all samples. Then, in all other bootstrap runs, we match each changepoint to the closest run in the reference. We found that location of the changepoints is consistent across bootstraps: on average, changepoints are located 0.093 CCF apart from the closest reference changepoint.

#### 3.3.3 Signatures with most changing activities

As shown by Figure 6, samples typically have only two or three signatures with high activities. These signatures are usually the most variable (up to 87.2% max change, 12% on average). Other signature have low activity and remain constant. On average 3.6% of overall activity is made up of low-activity signatures (with activity <5%). Low-activity signatures most likely appear due to the uncertainty of our signature activity estimates. As mentioned in section 3.3.2, a mean standard deviation of signature activities of 2.9%, thus, we remove signatures with activity less than 5% as they are within two standard deviations of 0%.

**Figure 6.**
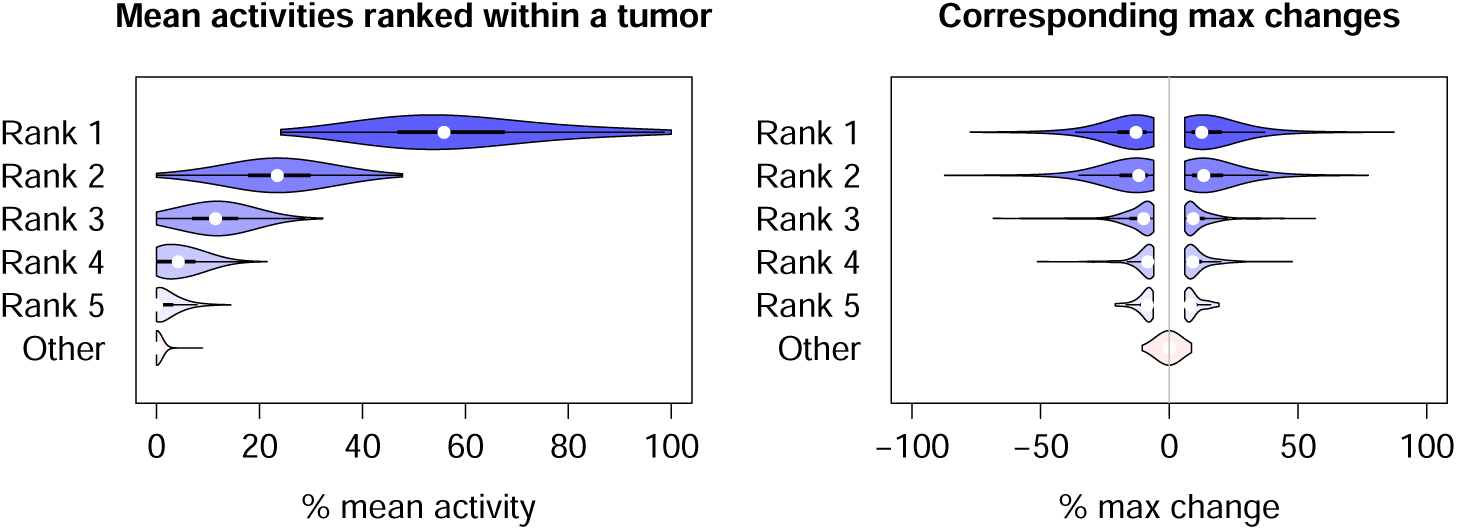
(a) Mean signature activities ranked from the largest to the smallest within each sample in PCAWG data. Only the top five signatures with the highest activities in a sample are shown. **(b)** Maximum changes of signature activities for the corresponding signatures on the left plot. The changes below 5% are omitted.

#### 3.3.4 Trends in signature change per cancer type

The majority of PCAWG samples have a signature change: 76.1% of samples have a max change >5% in at least one signature; 48.4% of samples have change >10%. However, the number of signature changes correlates to some extent with the number of mutations in the sample. Out of samples with less than 10 timepoints only 26.3% of samples have a change >5% compared to 80.4% across the rest of the samples (see distribution on fig. 7).

**Figure 7.**
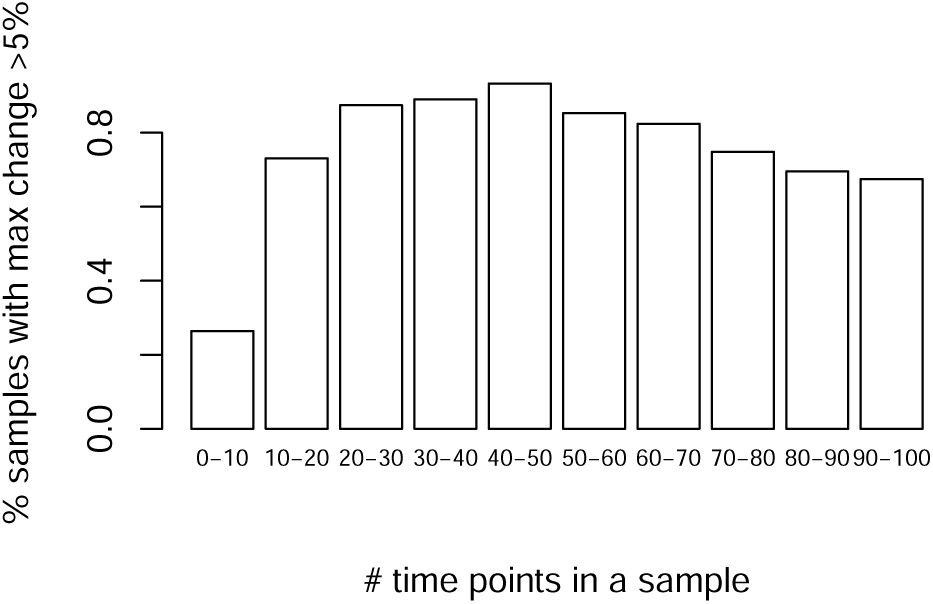
Proportion of tumours that have a significant change greater than 5% activity depending on the number of time points in a sample. Each bar corresponds to the range of number of time points in a sample.

### 3.4 Choosing active signatures

Only a subset of signatures are active in a particular sample, and this subset is largely determined by a cancer type. For the analyses reported above, we use a set of active signatures provided by PCAWG^36^, which contains a list of active signatures per sample (on average, four per sample). Frequent active signatures for each cancer type are available from a variety of sources^2, 36^.We highly recommend using either these cancer-specific active signatures or deriving sample-specific active signatures using one of the procedures described in this section. We strongly discourage using TrackSig with a full set of signatures on a single sample, as many of the signatures overlap considerably, which can cause signature activity estimation errors due to this collinearity.

Here we evaluate three different ways to select the active signatures, all supported by TrackSig. The first strategy, *all-sigs*, simply computes activity trajectories for all signatures. The second, *cancer-type-specific-sigs*, uses all signatures reported as active in the cancer type under consideration. The final strategy, *sample-specific-sigs*, first fits signature activities to the full set of mutation counts using an initial set of signatures, and sets the active signatures to be those with activities greater than a threshold (by default, 5%) in the initial fit. Then TrackSig computes activity trajectories only for the active signatures. In the following, we evaluate *sample-specific* when the initial set is *all-sigs*, however, we suspect this approach will also work well with *cancer-type-specific* as the initial set. We evaluate each strategy by comparing the active signatures selected by TrackSig with those reported by PCAWG-Signature group on the PCAWG tumour set^36^.

For *all-sigs*, we used all 48 signatures and we found on average, 44.7% of overall activity assigned by TrackSig is assigned to the active signatures selected by PCAWG-Signature group. Each incorrect signature gets 1.3% of activity on average. In other words, the incorrect activity is widely distributed among the signatures. Using *cancer-type-specific-sigs* improves the correspondence to 68.7% of the total activity on average. This strategy reduces the initial set of potentially active signatures from 48 down to 12 on average (ranging from 4 signatures in Lower Grade Glioma to 24 signatures in Liver Cancer). Here, we observe that signature 5 and 40 are the most prevalent among the incorrect signatures, having the average activity of 14% and 12.6% respectively in the samples where they are supposed to be inactive. Finally, if we use the *sample-specific-sigs* strategy starting with *all-sigs* as the initial set, we exactly recover the active signatures reported by PCAWG-Signature group.

Fitting either per-cancer or per-sample signatures results in more activity mass to be on the correct signatures and speeds up the computations. Therefore, we recommend choosing one of these instead of using activities from the full set.

## 4 Summary and Conclusions

TrackSig reconstructs the evolutionary trajectories of mutational signature activities by sorting point mutations according to their inferred CCF and then partitioning this sorted list into groups of mutations with constant signature activities. TrackSig estimates uncertainty in the location of the changepoints using bootstrap. TrackSig is designed to be applied to VAF data on SNVs from a single sample, however, it can be applied to either sorted lists of point mutations derived from subclonal reconstruction algorithms, or CCFs from a single cancer sample derived from methods which perform multi-sample reconstructions or subclonal CNA reconstructions.

Changepoints often correspond to boundaries between subclones^24^. In our simulations we show that TrackSig often better detects subclones than methods explicitly designed to find subclones, especially when there is a mismatch between the assumed and actual VAF generation process. By reconstructing changes in signature activities, TrackSig can potentially help identify DNA damage repair processes disrupted in the cell and, in doing so, help inform treatment^11^.

### 4.1 Relationship to previous work

Previous approaches estimate signature activities for a group of mutations without considering their timing (e.g. eMu^33^ or deconstructSigs^13^). Therefore, the attempts to compare activity changes across evolutionary history have relied on pre-defined groups of mutations, such as those occurring before or after whole genome duplications^7, 9, 29, 37^; those classified as clonal or subclonal^1, 9^; or those grouped in subclones via multi-region sequencing^15–19^. As such, the accuracy of these methods relies on i) the accuracy in grouping mutations based on VAF – which is low with data from a single bulk sample^21^; and ii) the existence of a small number of subclones or mutation groups within a sample, which is not true for neutrally evolving tumours^23, 25, 38^.

In contrast, TrackSig uses the distributions of mutation types to group mutations, this permits more accurate reconstruction of signature activities than clustering mutations by VAF alone. Indeed, as our simulations demonstrate, not only are the signature activities more accurately reconstructed, but in some cases, TrackSig is a more sensitive detector of subclones. Furthermore, TrackSig makes fewer assumptions about the underlying VAF distribution, so it can be readily applied to data from neutrally evolving tumour populations^25, 38^. Our simulations further demonstrate that TrackSig’s reconstructions are less sensitive to model misspecification errors, such as violations of the infinite sites assumptions.

Clustering methods applied to VAFs from single bulk samples require high read depth for accuracy^21^. Indeed, due to this challenge, previous approaches have used multi-region sequencing^15–^19, 23^, 38–41^. In contrast, TrackSig can be deployed in a much larger range of settings. Separately, we report that TrackSig can detect subclones that are missed by VAF clustering methods.^24^

Another important innovation of TrackSig is the use of CCF as a surrogate for evolutionary timing. Similar ideas have been used in human population genetics, where variant allele frequency to get relative order of mutations along the ancestral lineage^42^. In population genetics, allele frequency is calculated across individuals, while we calculate VAF across cell population within a single sample. In TrackSig we estimate cancer cell fraction (CCF) and reconstructions of clonal CNAs.

### 4.2 Sensitivity to misorderings of the SNVs

For ease of presentation, we have assumed that ordering SNVs by CCF recovers the order in which they accumulated in the genomes of ancestral cells. However, this assumption is not critical for correct reconstruction of signature activity changes.

First, we have shown through bootstrap sampling and the clonal evolution simulations that errors in the estimation of SNV CCFs due to sampling noise have a limited impact on TrackSig’s ability to estimate accurate activity trajectories. We have similarly shown that these activity trajectories are not impacted if a small fraction (3%) of the SNVs violate the infinite sites assumption.

However, these trajectories can be impacted by incorrect ordering of a large numbers of SNVs. These can occur in two ways. First, misordering can occur if a CNA changes the number of SNV allele’s per cell. For example, daughter cells can fail to inherit SNVs in their mother cells due to a loss of heterozygosity (LOH). If a CNA reconstruction is available, TrackSig will correct for any detected clonal LOH when ordering SNVs, and will not attempt to order SNVs in regions affected by subclonal CNAs, thereby resolving this difficulty. However, if a CNA reconstruction is not available, or it is inaccurate, the accuracy of the activity trajectories can suffer. As such, we recommend only using TrackSig when CNA reconstructions are available and reliable.

Second, SNV ordering need not correspond to time of acquisition when a single sample contains SNVs from subclones from different branches of the cancer phylogeny. In these circumstances, there is not a single linear order for the activities, and furthermore late occurring subclones on a different branch can have higher CCF than earlier ones occurring in the sample. This situation also occurs when the sample contains a large number of neutrally evolving mutations from multiple subclonal lineages. Note that such circumstances are rare in single biopsies^29^ and that furthermore, a subclone can only be misordered if its CCF is less than 50% due to the Pigeonhole Principle^1^, so the ordering by CCFs in guaranteed to be correct up until 50% CCF. However, even when these misorderings occurs, our simulations demonstrate that, with one exception, TrackSig’s activity reconstructions, and estimation of number of subclones, are largely unaffected.

Even in the rare circumstance that SNV misordering does occur, it may be possible to detect it, and interpret the activity changes correctly. For example, if late occurring but misordered SNVs manifest a more drastic change in signature activity, this misordering may be detectable by the presence of oscillations in the activity trajectories. To address this issue, when assessing overall change in signature activity, we computed the difference between the lowest and highest activities for each signature. This difference will be consistent regardless of ordering.

### 4.3 Applicability to other mutation types and multi-region sequencing

In TrackSig, the number of mutation types is provided as a parameter and is not fixed to 96 types. Because of this, it is straightforward to generalize TrackSig to reconstruct the activities of different mutation signatures or different mutations, so long as these mutations can be approximately ordered by their evolutionary time and each mutation can be classified into one of a fixed number of categories. In this paper, we ordered SNVs by decreasing CCF. This same strategy could be naturally extended to indels for which the infinite sites assumption is also valid. The infinite sites assumption should also be valid for structural variants (SVs) associated with well-defined breakpoints, thus permitting TrackSig to be used to track activities to recently defined SV signatures^37^. The CCFs of SVs can be estimated using the VAFs of split-reads mapping to their breakpoints^43^. Because they cover larger genomic regions, infinite sites is less valid for CNAs, although it is possible to approximately order clonal CNAs based on the inferred multiplicity of SNVs affected by them^9^.

TrackSig also requires a pre-defined set of mutation signatures, each of which is a probability distribution over the mutation types. However, if these signatures are unavailable, they can be defined by non-negative matrix factorization, or Latent Dirichlet Allocation^44^, if counts across mutation types are available from multiple cancer samples.

TrackSig can be applied to VAFs from bulk sequencing data from multi-region sequencing or longitudinal samples by simply running it on each sample separately. In preliminary experiments testing this approach we found broad consistency in the active signature selected, and in the signature activities of the clonal mutations in each sample. We observe with only 0.03% mean absolute activity difference (0.017 KL divergence) between signature activities of clonal cluster across different samples. See appendix E and Suppl Figure F.9 for details.

### 4.4 Accounting for changes in mutation rate

The timelines reconstructed by TrackSig are computed with a fixed number of mutations in each bin. If overall rate of generating mutations in tumour was constant, our timeline would correspond to the real time. However, tumour mutation rate often accelerates throughout development^45, 46^. Although the changing rate does not affect our analysis, the estimates of the pseudo-time might not be linearly related to real time.

Estimating changes in overall mutation rate is difficult. A possible way to correct for this is to adjust the time line based on activities of signatures 1 and 5. Some report that signatures 1 and 5 operate as a cell “clock” as the number of mutations contributed by these signatures is proportional to the age of the individual^6^. Determining the association between our pseudo-time estimates and real time is left for further investigation.

Our method TrackSig provides further insight how signature profile changes throughout tumour development. We show that through signatures analysis we can detect major events in tumour evolution, notably, transitions to a new subclone. Mutational signatures provide a unique way to recover tumour evolution path, track activities of mutational processes, adjust the treatment strategy and detect changes in therapy response.

## Author’s contributions

QDM designed the project and supervised the study. YR designed and implemented the method and performed the experiments. YR and QDM wrote the manuscript with assistance from CH and RS. YR and CH made figures. RS implemented PELT algorithm. RL performed the non-parametric simulations. CH and YR performed clonal evolution simulations. CH implemented the SciClone+DeconsructSigs baseline. JW and AD provided assistance with tumour phylogeny reconstruction. NL wrote the script to extract tri-nucleotide counts. The PCAWG network provided the mutational signatures and feedback on the method. All authors read and approved the final manuscript.

## Availability of data and materials

Code is available at https://github.com/morrislab/TrackSig. Code for generating simulations is included in the Github repository.

## Acknowledgments

We thank Pan-cancer Analysis of Whole Genomes (PCAWG) network, and in particular the PCAWG Evolution and Heterogeneity working group, for providing data, analysis and valuable input on this project. We would in particular like to highlight Peter Van Loo, Clemency Jolly, Stefan Dentro, David Wedge, Paul Boutros, Lydia Liu, and Moritz Gerstung who provided valuable feedback during the development of the TrackSig methodology. We would like to acknowledge SciNet as part of Compute Canada for providing computational resources. This research was partially supported by an Natural Science and Engineering Research Council operating grant; an Associate Investigator award from the Ontario Institute of Cancer Research; and a subgrant from the Canadian Centre for Computational Genomics genomics technology platform funded by Genome Canada, all to QDM. It also received funding from the University of Toronto’s Medicine by Design initiative, which in part of the Canada First Research Excellence Fund (CFREF) and the Compute the Cure gift from the NVIDIA foundation. QDM is a Canada CIFAR AI chair at the Vector Institute.

## A Computing activity to mutational signatures

We apply topic modeling^47^ to infer signature activities. Within the time point, we separate mutation into K mutation types. Mutation types relate to vocabulary in topic modeling. The types used in TrackSig are described in section 3.1. Then we use mixture of discrete distributions to infer signature activities. We describe this model below.

We represent each mutation as a *K*-dimensional binary vector – “one-hot-encoding” of a mutation type. “One-hot-encoding” of a mutation of type *k* is a binary vector where *k*-th component is equal to 1, and other components are zeros. We will denote **x**^(*n*)^ to be the “one-hot-encoding of mutation *n*. A sample containing *N* mutations is represented as a *N × K* binary matrix **X**, where each column corresponds one mutation.

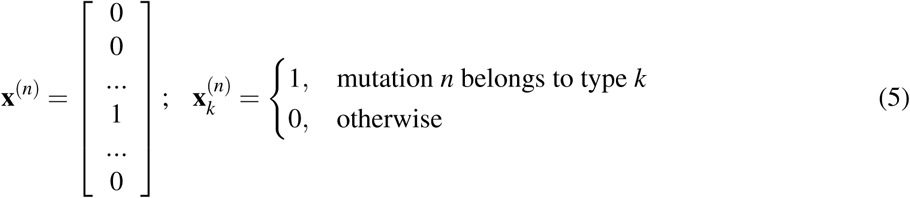

A mutation process is represented as a distribution over mutation types, known as a “*mutation signature*”. We will denote signature multinomials as *K*-dimensional probability vectors ***µ***_*i*_, where *i* = {1..*M*}is an index over signatures. Signatures are fixed and are not updated during the training.

We aim to estimate signature *activities* ***π*** – the proportion of mutations generated by each signature.

We will use the following notation:

*K* – number of mutation types

*M* – number of signatures

*N* – number of mutations

**x**^(*n*)^ – *K*-dimensional binary vector of mutation *n*

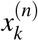 – *k*-th component of vector **x**^(*n*)^

***µ*** _***i***_ – i-th signature (*K*-dimensional vector)

*µ*_*ik*_ – k-th component of vector ***µ***_***i***_

***π*** – signature activities (mixture coefficients, *M*-dimensional vector)

*π*_*i*_ – i-th component of ***π*** (signature activity of signature *i*)

*z*_*n*_ – signature assignment for mutation *n*

We represent mutation matrix **X** as a mixture of signature multinomials ***µ***_**1**_,..***µ _K_*** with mixture coefficients ***π***:

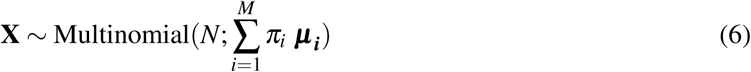

We denote *z*_*n*_ to be the signature assignment of mutation *n*. The probabilities of mutation *n* to be assigned to *i*-th signature are equal to the mixing coefficients:

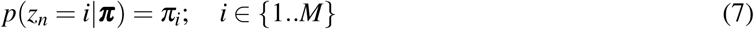

The probability of a mutation *n* to be generated by signature *i* is given by:

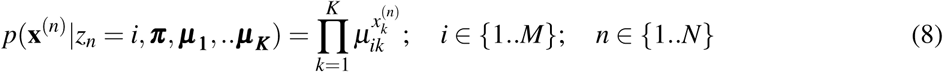

Then log likelihood of the collection of mutations in a sample:

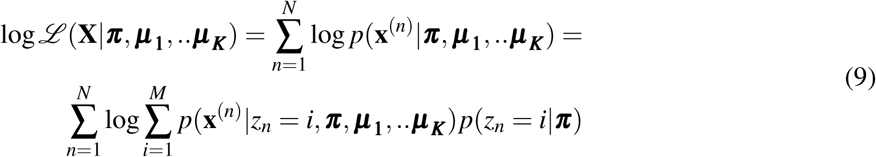

To estimate the activities, we fit mixing coefficients ***π*** in each bin using Expectation-Maximization (EM) algorithm^32^. The EM algorithm iterates between updating a posterior distribution over *z*_*n*_ and updating an estimate of the mixing coefficients ***π***

We start with initializing EM algorithm with uniform mixing coefficients:

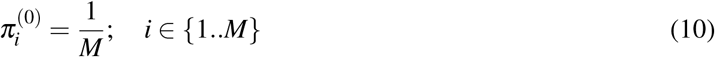

Then, we repeat the following E-step and M-step until the algorithm converges.

In E-step, at the *t*-th iteration, the posterior probabilities of mutation assignments to signatures are estimated as such:

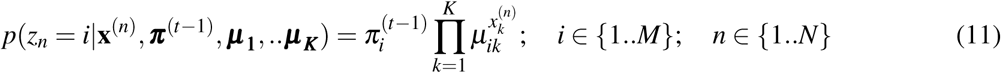

In M-step we update the estimates of the mixing coefficients:

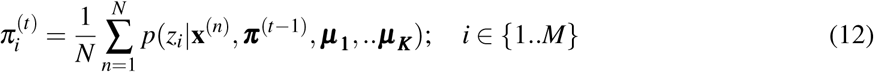

The algorithm has converged when the value of *π* is updated by less than 0.001 between iterations. The resulting mixture coefficients as the activities of the mutational signatures. We show the activities as percentage for the convenience of interpretation.

## B Pruned Exact Linear Time (PELT) Algorithm

We adapt Pruned Linear Exact Time (PELT)^34^ algorithm to detect change points in activity trajectories given cost function (likelihood) and BIC penalty. PELT is based on dynamic programming and uses heuristics to prune the set potential changepoints, thus reducing the computational time.

In this section, we will use the following notation:

*T* – number of time points

*P* – number of changepoints

*M* – number of signatures

### B.1 Locating change points

As described in 2.1.2, we separate mutations into bins 100 mutations, each of which represents one time point. Our input is the set of mutation counts across 96 types for each time point: *y*_1:*T*_ = (*y*_1_, …, *y*_*T*_). We aim to find *P* changepoints, or in other words, *P* + 1 segments. We denote *τ*_1:*P*_ = (*τ*_1_, …, *τ*_*P*_) to be the boundaries for our segments, meaning each segment will contain the data points 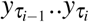.

Given a set of changepoints we can compute the likelihood of the data the following way. We fit mutational signatures within each segment (treating all mutations within each segment as one bin) and compute the likelihood 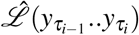 as described in A. The total likelihood is the sum of likelihoods in each segment:

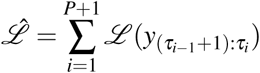

We aim to minimize the Bayesian Information Criterion (BIC):

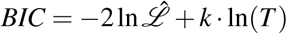

where *k* is the number of parameters in our model and *T* is the number of time points. In our case *k* = (*P* + 1)·(*M* −1) as we fit (*M* −1) signature activities in (*P* + 1) segments (recall that signature activities sum to 1).

We adapt PELT objective to minimize the BIC criterion. PELT aims to minimize sum of cost functions at each time point, while using a penalty *β* for each placed changepoint

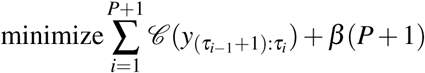

Intuitively, we are trying to select changepoints which result in the lowest cost (or highest likelihood) while reducing the penalty associated with adding changepoints. We set the parameters as follows to make the PELT equivalent to BIC:

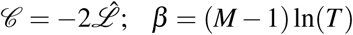

TrackSig-PELT algorithm finds the changepoints as follows. The algorithm starts with finding a partial solution in a subset of the timeline and then increases the search space until changepoints are located over the whole timeline. The algorithm keeps track of the time points *R*_*τ*_* that satisfy the pruning condition and which will be considered as potential changepoints at further iterations. At each iteration *τ*\*, the algorithm considers adding a new changepoint out of the set of available time points *R*_*τ*_*. To score a potential new changepoint, the algorithm refits the activities in bins formed by a potential changepoint. It finds a time point *τ′* with the smallest likelihood and adds it to the list of changepoints *cp*. Then the list of available time points *R*_*τ*_ * is updated: the potential changepoints are removed from further consideration if the increase in likelihood associated with this changepoint does not exceed the complexity penalty *β*.

### B.2 Pruning

PELT provides an improvement in runtime by pruning certain changepoints from consideration. We prune time point *t* if for all *t* < *s* < *T*:

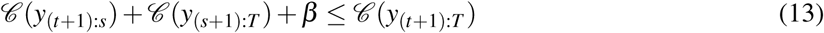

The cost of placing the last changepoint prior to *T* at *t* will always be higher than cost of placing the last changepoint prior to *T* at *s*. Given this result, we can eliminate *t* as a potential changepoint for all iterations of the dynamic programming algorithm as it will never be optimal going forwards.

#### Algorithm 1 TrackSig PELT Method (Killack and Eckley 2012)

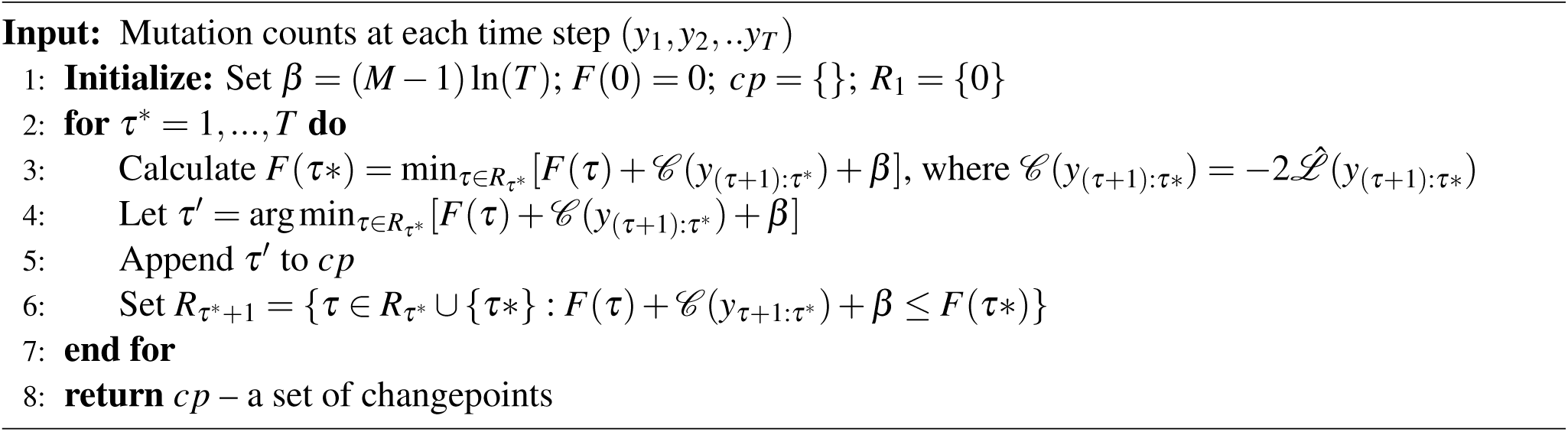

## C Clonal evolution simulations

### Choice of signatures

We generate the simulations with four active signatures: S1, S5 and two randomly-sampled signatures, which we will call A1, A2. Two other signatures A1, A2 are sampled from uniformly from the set of PCAWG (excluding signatures S1, S5, S7 and “artifact signatures” S40-S60). We decided to exclude signature S7 (sum of signatures S7a, S7b, S7c, S7d) as it had a distribution similar to uniform and was easily confused with other signatures both by TrackSig and DeconstructSigs. We include signatures S1 and S5 in all simulations as they are present in all real samples in PCAWG.

We sample activities separately for each cluster. We sample the activity of S1 from [0.03, 0.1] interval, S5 from [0.05, 0.15] interval, A1 from [0.4, 0.7] interval. The remaining activity is assigned to signature A2 (all signature activities have to sum to 1).

### Sampling mutation types

To sample mutation types from a signature, we treat it as a multinomial distribution and sample from it. The number of mutations sampled from each signature is equal to the activity of this signature multiplied by the total number of mutations.

### Sampling number of ref and alt alleles

Here we describe sampling number of ref and alt alleles for each mutation of the cluster, given the cluster CCF, number of mutation in the cluster and desired mean mutation depth. We tested mean mutation depths of 10, 30 and 100.

For each mutation, we sample read depth *d* from Poisson distribution with specified mean depth. Then we compute the probability of alt allele as 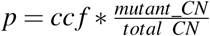, where *cc f* is CCF of the current cluster. Finally, we sample number of alt alleles *a* from a *Binomial*(*d, p*) and set the number of ref alleles to be the difference between depth and alt alleles.

In simulations with one and two clusters we use normal copy number of 2, mutant copy number of 1 and purity 1. Each simulation has 5000 mutations in total. We generate 100 simulations of each of five simulation types (one-cluster, two-cluster, branching, cna gain and infinite site assumption) and for each read depth that we tested.

### C.1 Basic simulations

First, we create simple one- and two-cluster simulations.

#### One-cluster simulations

We create one cluster with the average cluster CCF=1. Number of ref and alt alleles for each mutation is sampled as described in the previous section. We sample activity of the first active signature A1 from the interval *Uniform*([0.4, 0.7]), activity of time-related signature S1 from *Uniform*([0.03, 0.1]), and time-related signature S5 from *Uniform*([0.05, 0.15]). The remaining activity is assigned to the signature A2. Finally, we sample mutation types from each of active signatures. Number of mutation types sampled from each signature is proportional to their activities.

#### Two-cluster simulations

We create the first cluster with CCF=1 as described above. For the second cluster we sample ccf from *Uniform*([0.2, 0.6]) distribution. To sample signature activities, we follow the produre similar to one-cluster simulations. We sample activity of the first active signature A1 from *Uniform*([0.4, 0.7]) for the clonal cluster, and *Uniform*([0.2, 0.4]) for the second cluster to ensure the signature activity change between the two clusters. Full procedure is shown in Algorithm 2.

### C.2 Branching

To test violation of TrackSig assumptions, we create simulations with branching, CNA gain or violation of infinite site assumption. To simulate branching, we create three clusters. The clonal cluster is always assigned CCF=1. The CCF for the last cluster (with the smallest CCF) is sampled uniformly from [0.2, 0.35]. The middle cluster CCF is sampled such that it has at least 0.15 gap on CCF scale with other clusters. Additionally, we ensure that sum of CCFs of the second and third clusters does not exceed 1 (otherwise the clusters cannot be branched).

In branching simulations, we expect to see the signature activity for A1 signature decreases at the transition to the second cluster and increases again at the transition to the third subclone. If such step-like behavior of is observed in real data, we consider this a sign of branching. Note that if we reversed the order of the branched clusters and assigned the same signature activities to the first and second clusters, it woudn’t be possible to distinguish between these two clusters since TrackSig can only find changepoints based on signature change.

To show the effect of branching on signature trajectories, we assign similar activities to the first (clonal) and third cluster (with the smallest CCF), but introduce a signature change in the second (middle) cluster. To do this, we sample signature activity for A1 from Uniform([0.4,0.7]), calculate the exposures for other signatures and assign the same activities to the first and last cluster. For the middle cluster, we sample activity for A1 signature from Uniform([0.2, 0.4]). As before, we sample activity of time-related signature S1 from *Uniform*([0.03, 0.1]), and time-related signature S5 from *Uniform*([0.05, 0.15]) and assign the remaining activity to A2.

### C.3 CNA gain

CNA gain simulations are based on the branching simulations described above and has three clusters: clonal and two subclones.

We introduce a CNA gain for 10% of mutations in the clonal cluster: 5% of mutations have CNA gain on the mutant allele and 5% have CNA gain on reference allele. Thus, 10% of mutations get total copy number 3 and mutant copy number of 2 and 1 respectively. We assume that these copy number changes are inherited by both subclones. To simulate the CNA change, we adjust the *mutant*_*CN* and *total*_*CN* parameter in Algorithm 3 for 10% of mutations in each cluster. We provide total copy number a input to both TrackSig and SciClone.

### C.4 Violation of infinite site assumption

To simulate the violation of infinite site assumption (ISA), we create four clusters. The first three clusters are created the same way as in the branching simulation. The forth cluster simulates mutations that occurred in both clusters independently, thus violating ISA. The CCF of the forth cluster is the sum of CCFs of the two subclonal clusters. We assign 3% of all mutation to the forth cluster. As expected, the presence of mutations that violate ISA don’t affect signature activity trajectories.

### C.5 Neutral Evolution Mutations

To make our simulations more realistic, we add mutations which emerged due to neutral evolution. We follow Williams et al. ^25^ for generating mutations from neutral evolution. First, we establish the number of neutral mutations to be generated. Then we sample those mutations according to the power-law distribution 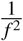, where *f* is variant allele frequency. Both steps are described in more detail below.

The number of neutral mutations is computed as follows:

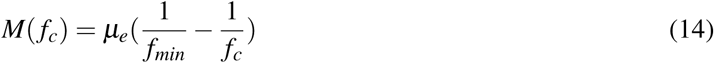

where *f*_*c*_ is the variant allele frequency (VAF) of the cluster, *f*_*min*_ is a minimal VAF in consideration and *µ*_*e*_ is effective mutation rate. For clonal cluster, *f*_*c*_ = 0.5. We only consider mutations with 3 or more mutant reads. Therefore, we set 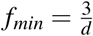, where *d* is the mean depth of the simulation. We use *µ*_*e*_ = 16^27^.

Next, we sample *M*(*f*_*c*_) mutations according to the power-law distribution on interval [*f*_*min*_; *f*_*c*_]. Cumulative distribution function (CDF) of power-law distribution on the interval [*f*_*min*_; *f*_*c*_] is the following:

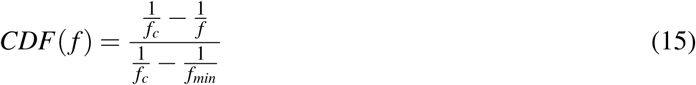

To sample from this distribution, we take samples from uniform distribution and then use inverse cumulative distribution function (I-CDF) to transform them into samples from power-law distribution. Inverse CDF function takes the following form:

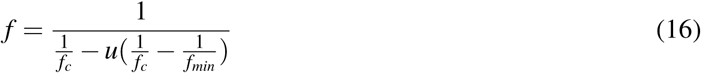

where *f* is our target allele frequency (i.e. sample from the power-law) and *u* is a sample from uniform distribution.

After discussion with our colleagues from population genetics, we decided to generate neutral mutations only from a clonal cluster.

Our results are shown in additional bar in figures 3b and F.2. At depth 10 and 30 TrackSig’s ability to detect subclones is not impacted by neutral mutations. At depth 100, both TrackSig and SciClone detect an extra cluster, which is consistent findings of Williams et al.^25^: neutral evolution can be detected at a minimal depth of 100. Figure 4b shows the example of generated simulation at depth 10.

## D SciClone+DeconstructSigs baseline

To showcase the potential of our method, we compared TrackSig to SciClone+DeconstructSigs pipeline which is commonly used to infer signature activities.

### D.1 SciClone + DeconstructSigs

First, we clustered SNVs using SciClone (v1.1)^35^. Sciclone uses variational Bayesian mixture model to cluster SNVs based on their CCF. We provided CNA calls as a part of input for SciClone, same as we do in TrackSig. Since we needed to test clustering at low depth, we used minimum read depth of 1. We report the results with two clustering methods in SciClone: Beta mixture model (BMM, default) and Beta-binomial mixture model (Binomial BMM).

Finally, we took the mutation clustering performed by SciClone and computed activities of mutational signatures withing each cluster using DeconstructSigs (v1.8.0)^13^. We used the same set of PCAWG signatures as we used in TrackSig. We fit the same set of active signatures with DeconstructSigs as we do in TrackSig.

### D.2 PyClone

We attempted to use PyClone (v0.13.1)^30^ instead of SciClone. However, PyClone uses a Markov Chain Monte Carlo (MCMC) approach, and has a time complexity of *O*(*n*2). This is feasible for the number of mutations validated on in the paper PyClone is described, but quickly becomes intractable for whole genome sequencing containing thousands of mutations. We didn’t manage to run the on samples containing more than 1000 SNVs.

## E Analysis of multi-region cases

To compare mutational signatures across multiple samples, we run TrackSig separately on each sample. Samples from the same tumour can have different active signatures. Therefore, for each tumour, we split the samples into groups that have the same set of active signatures and compare samples only within the group. To compare signatures of the clonal cluster, we compute KL divergence and mean activity difference between the first time points of the samples with the same active signatures. Within each group of samples with the same set of active signatures, we compute the pair-wise mean activity difference and KL difference between all pairs of samples within the group. We report the mean metrics of all pairs within the signature group.

### Algorithm 2 Simulation algorithm for two clusters

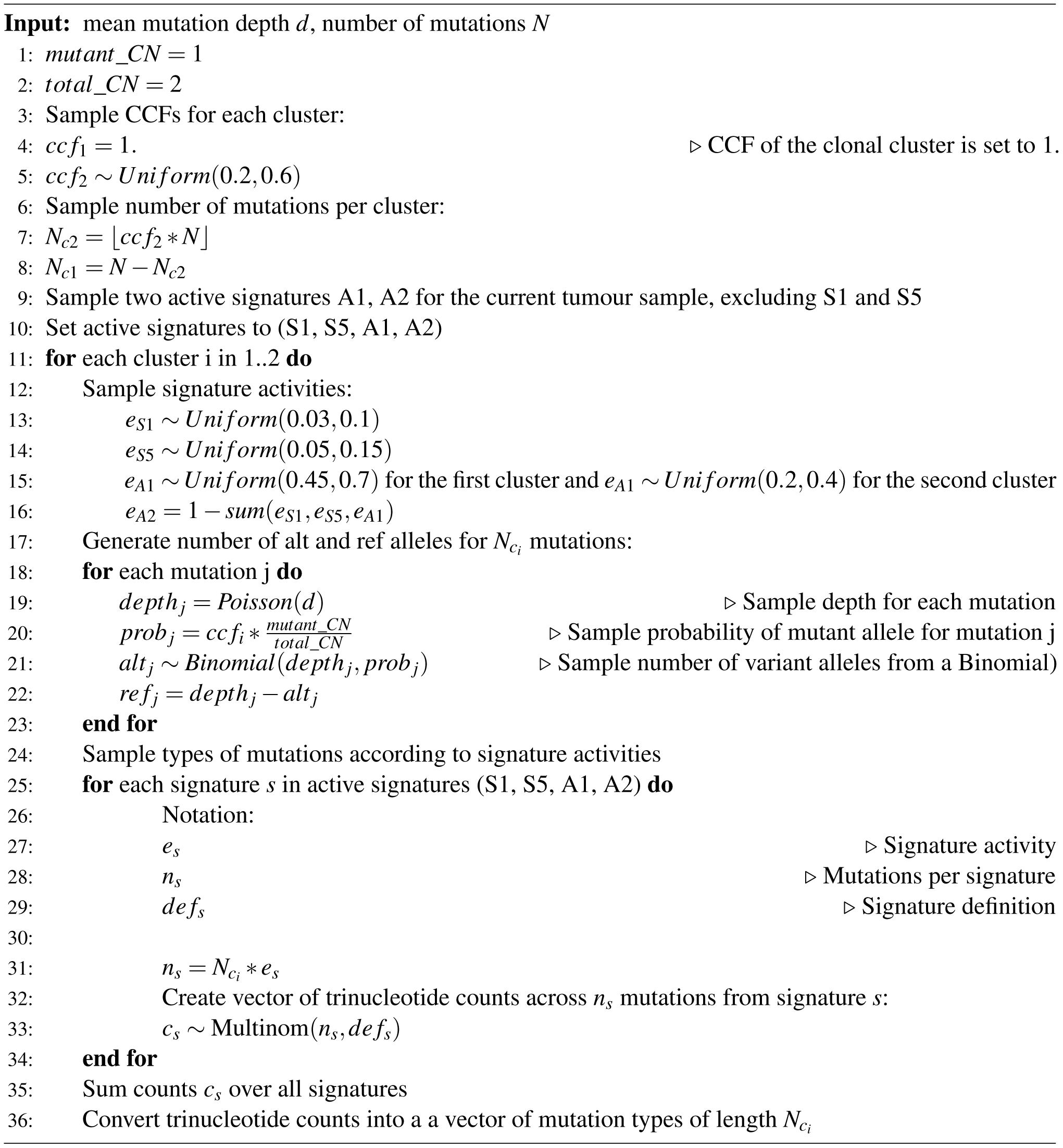

### Algorithm 3 Simulation algorithm for branching with three clusters

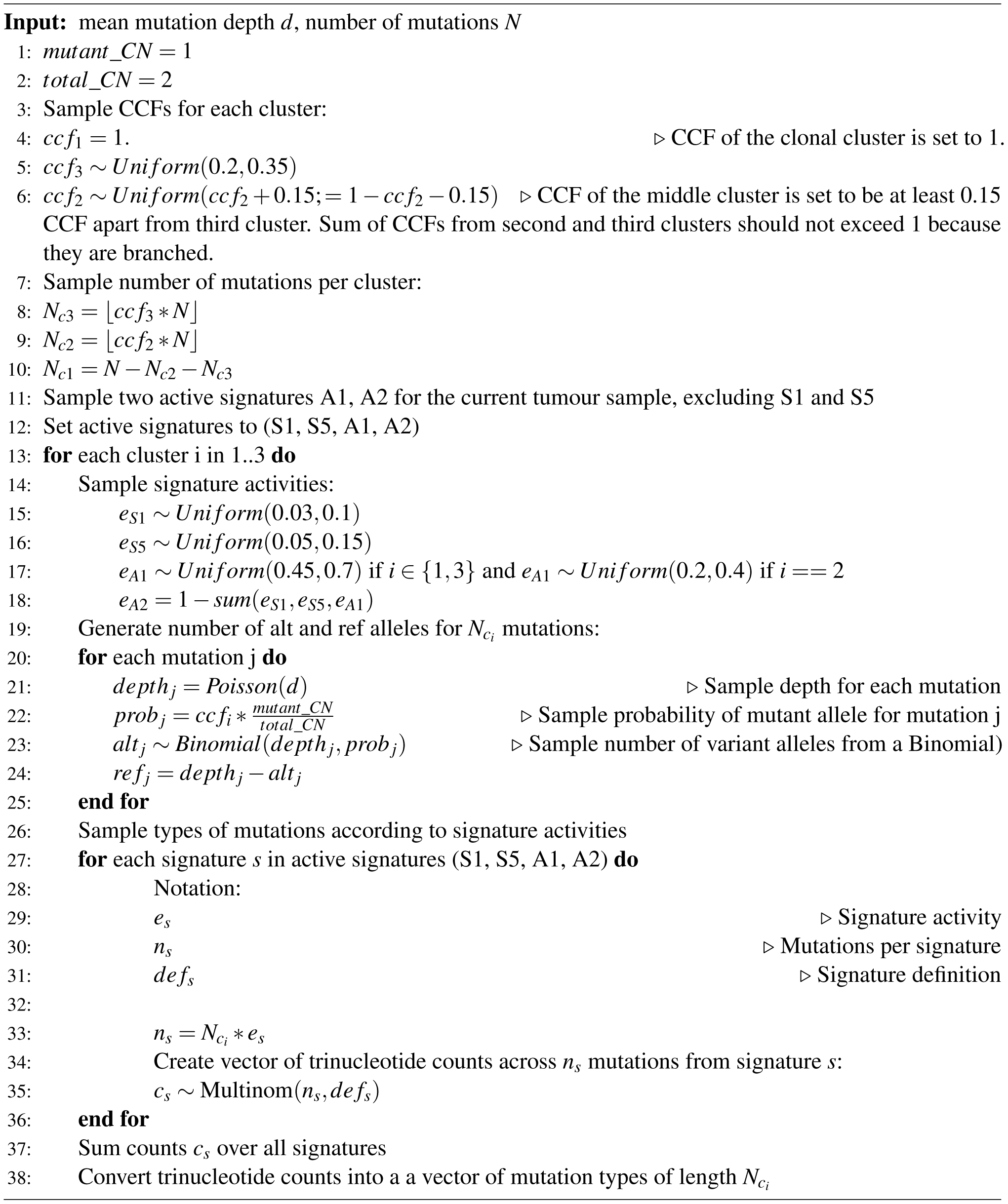

## F Supplementary figures and tables

**Figure F.1.**
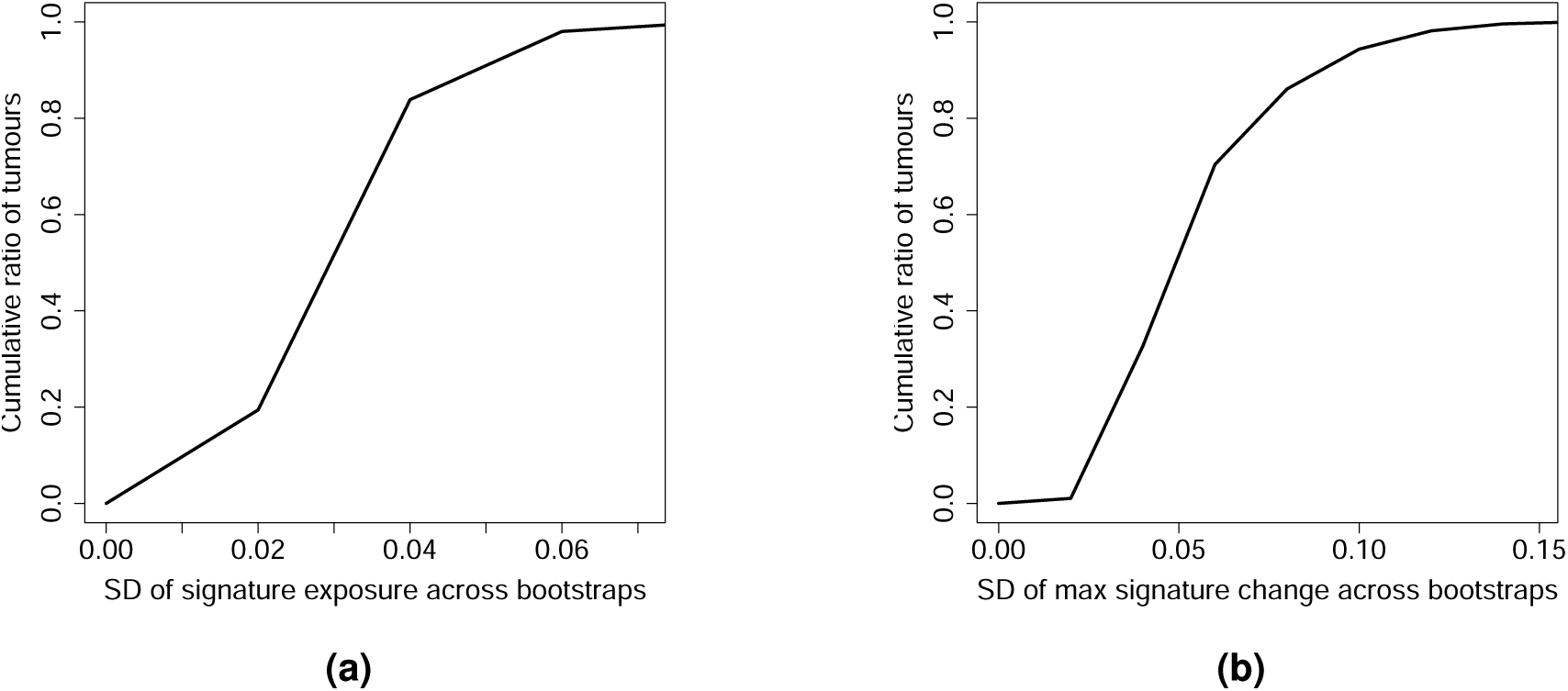
Discrepancies in signature activities on bootstrap data. **(a)** Standard deviations of signature activities at each time point for each signature across bootstraps. **(b)** Standard deviations of activity *change* at each time point for each signature across bootstraps.

**Figure F.2.**
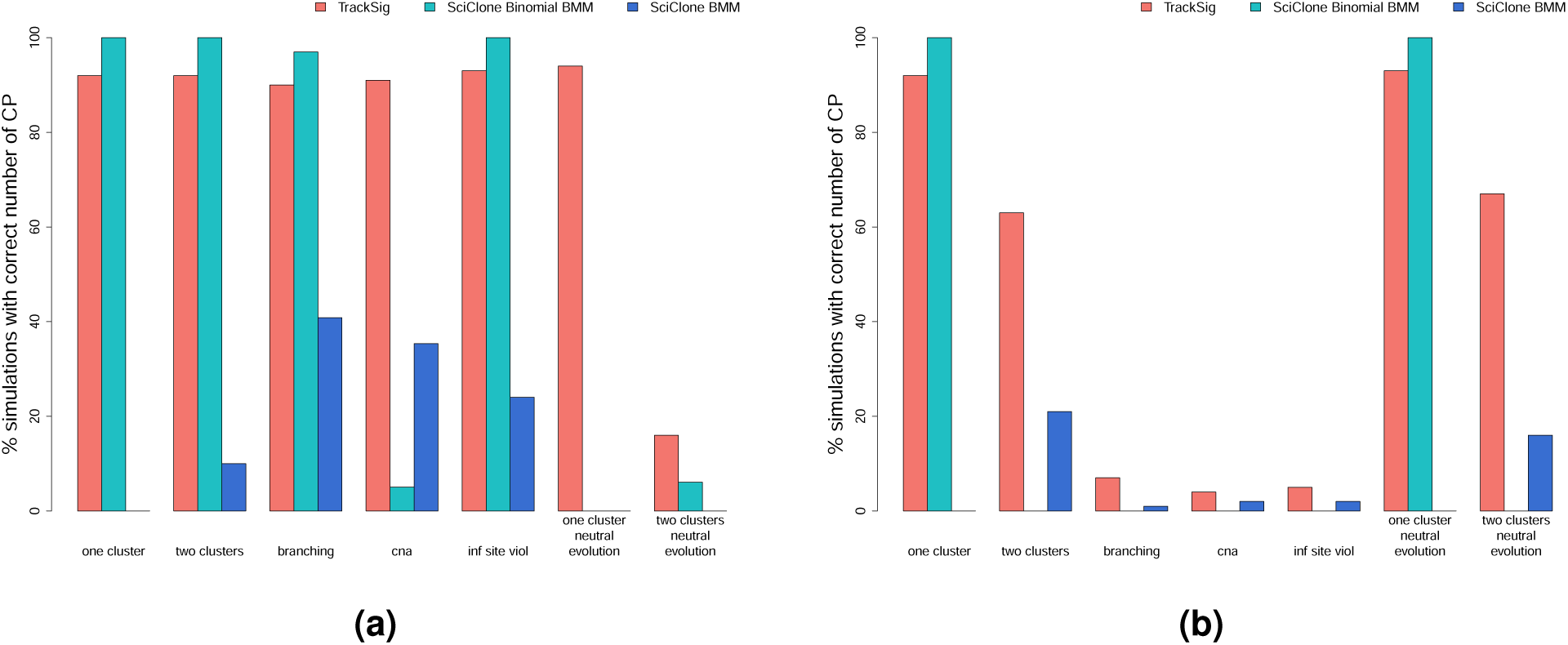
Comparison of TrackSig and SciClone for correctness in number of subclones detected. Each method was evaluated on all simulation scenarios in Appendix C, shown in X-axis. Y-axis shows the percentage of simulations where the method predicted the correct number of changepoints. Comparison was performed on simulated data with read depth **(a)** 100 and **(b)** 10.

**Figure F.3.**
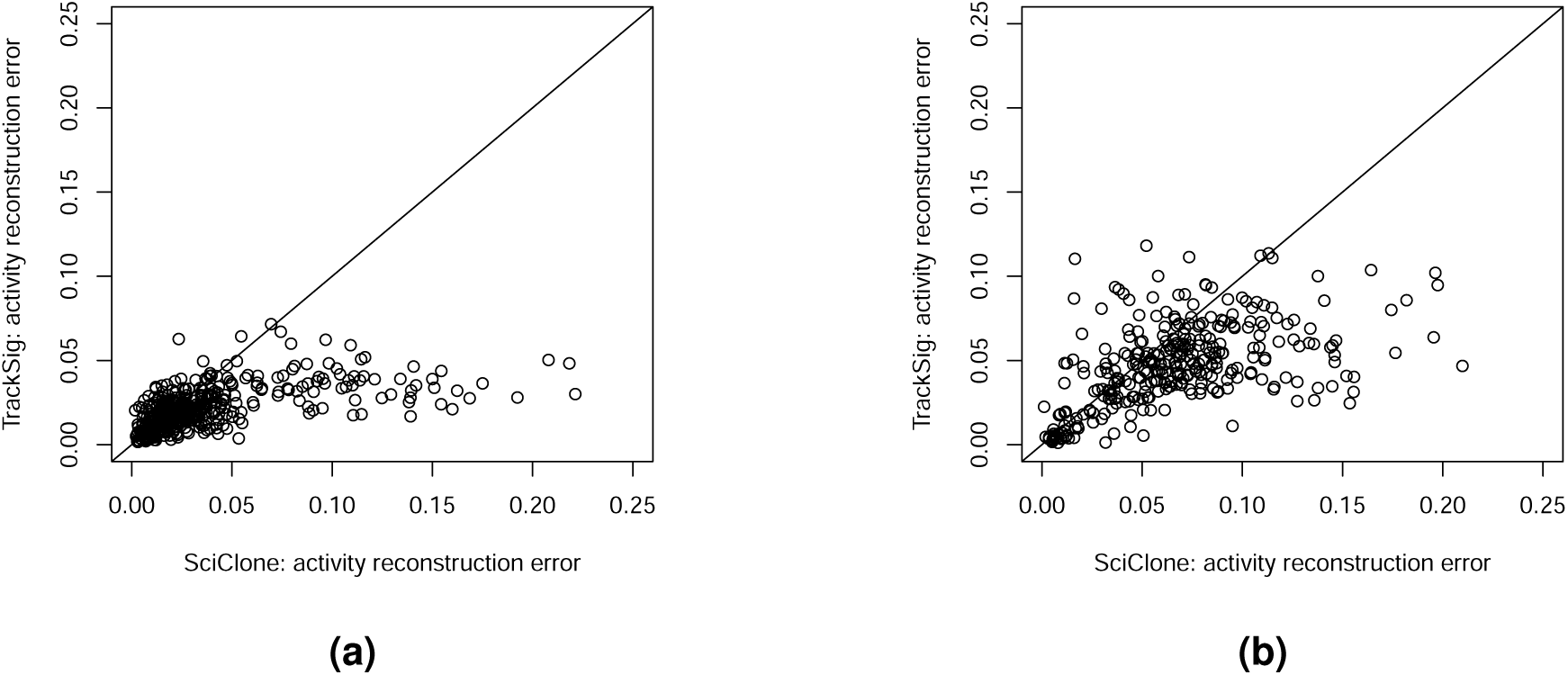
Median per-mutation activity reconstruction error (absolute activity difference) between the method (TrackSig and SciClone) and the true activities on clonal evolution simulations. **(a)** Depth 100. Mean activity error: TrackSig 0.022, SciClone 0.039. **(b)** Depth 10. Mean activity error: TrackSig 0.048, SciClone 0.068.

**Figure F.4.**
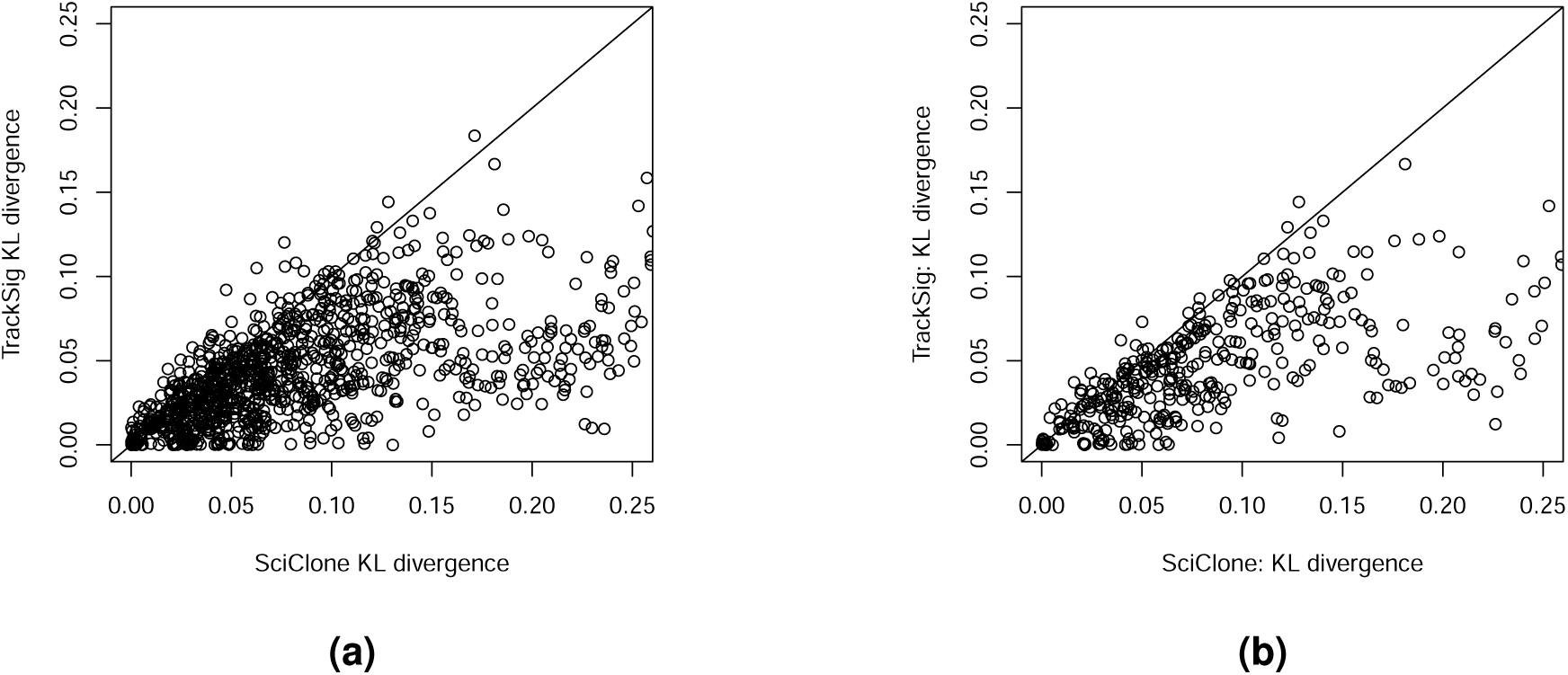
Mean per-mutation KL divergence between predicted and true exposures on clonal evolution simulations. **(a)** All simulations. Mean per-mutation KL divergence: TrackSig 0.044, SciClone 0.091. **(b)** Depth 30. Mean KL divergence: TrackSig 0.047, SciClone 0.095.

**Figure F.5.**
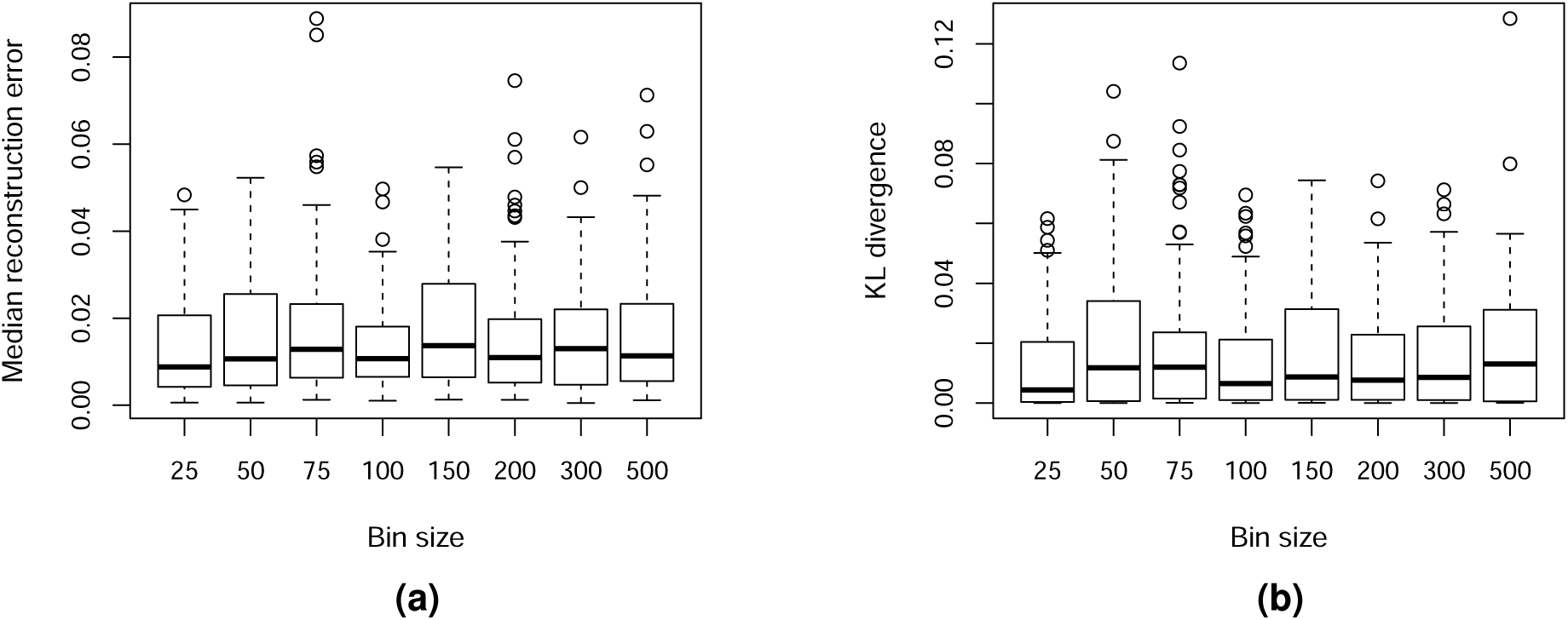
**(a)** Median absolute, per-mutation difference between true activities and activities estimated by TrackSig for different bin sizes at depth 30. **(b)** Mean per-mutation KL divergence between estimated and true activities for different bin sizes at depth 30.

**Figure F.6.**
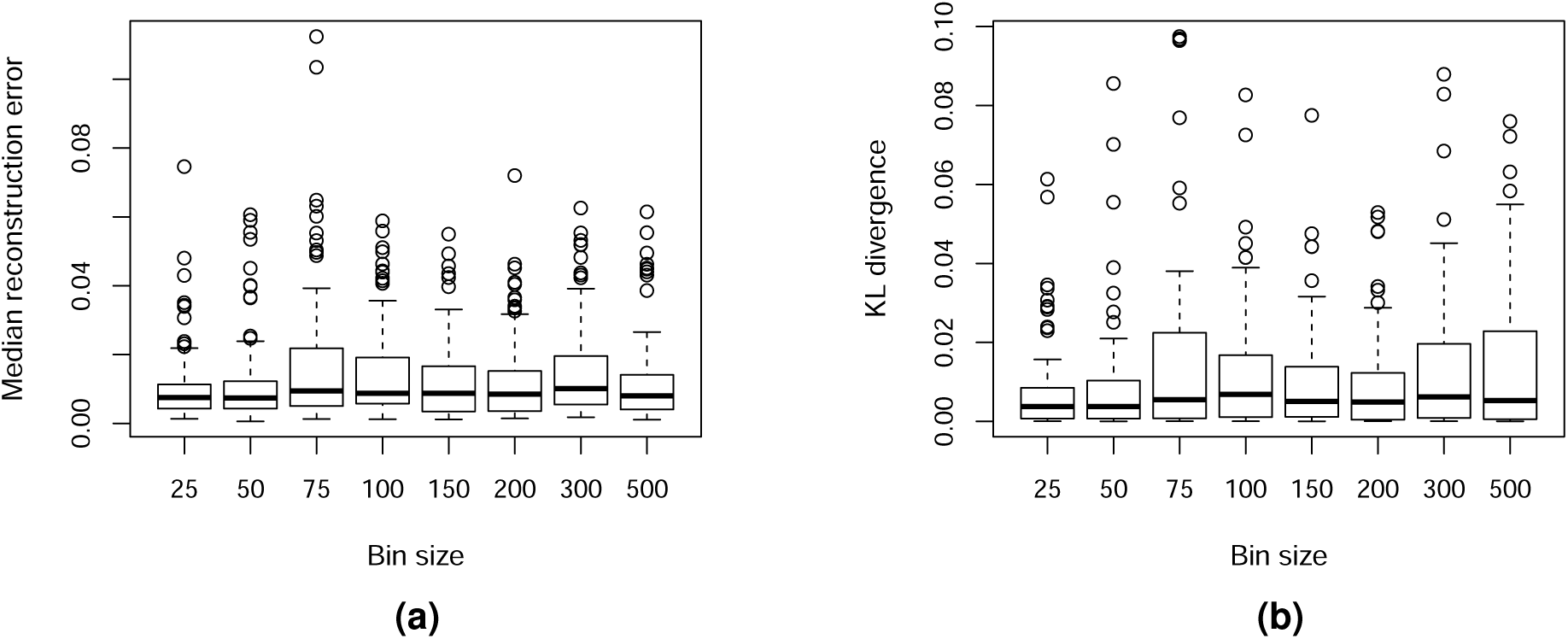
**(a)** Median absolute, per-mutation difference between true activities and activities estimated by TrackSig for different bin sizes at depth 100. **(b)** Mean per-mutation KL divergence between estimated and true activities for different bin sizes at depth 100.

**Figure F.7.**
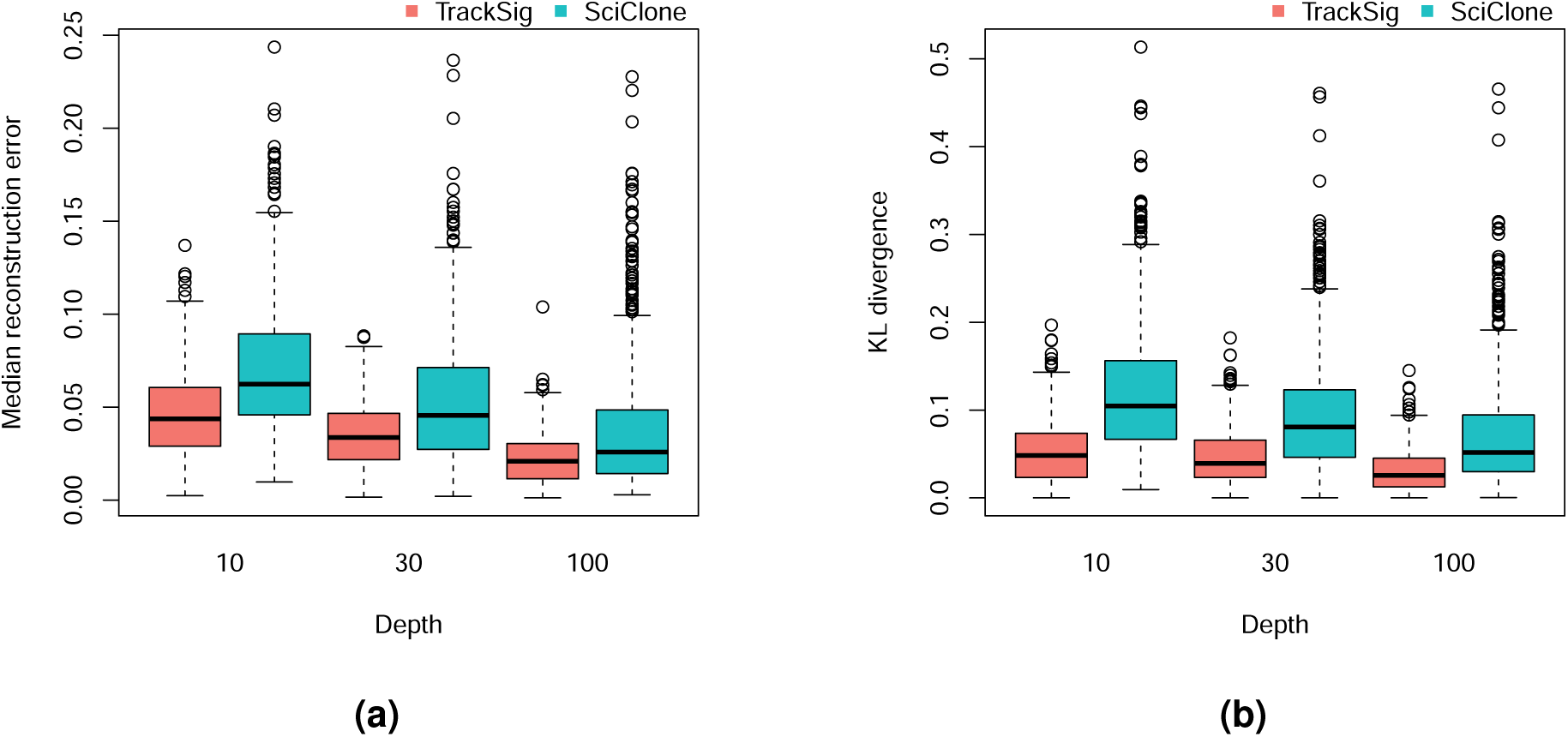
Comparison of TrackSig and SciClone (BMM noise model, default) on simulation scenarios described in Appendix appendices C.1 – C.4. Performance was evaluated across different simulated read depths, shown in X-axis. **(a)** Median absolute, per-mutation difference between true activities and activities estimated by each method. **(b)** Mean per-mutation KL divergence between true activities and activities estimated by each method.

**Figure F.8.**
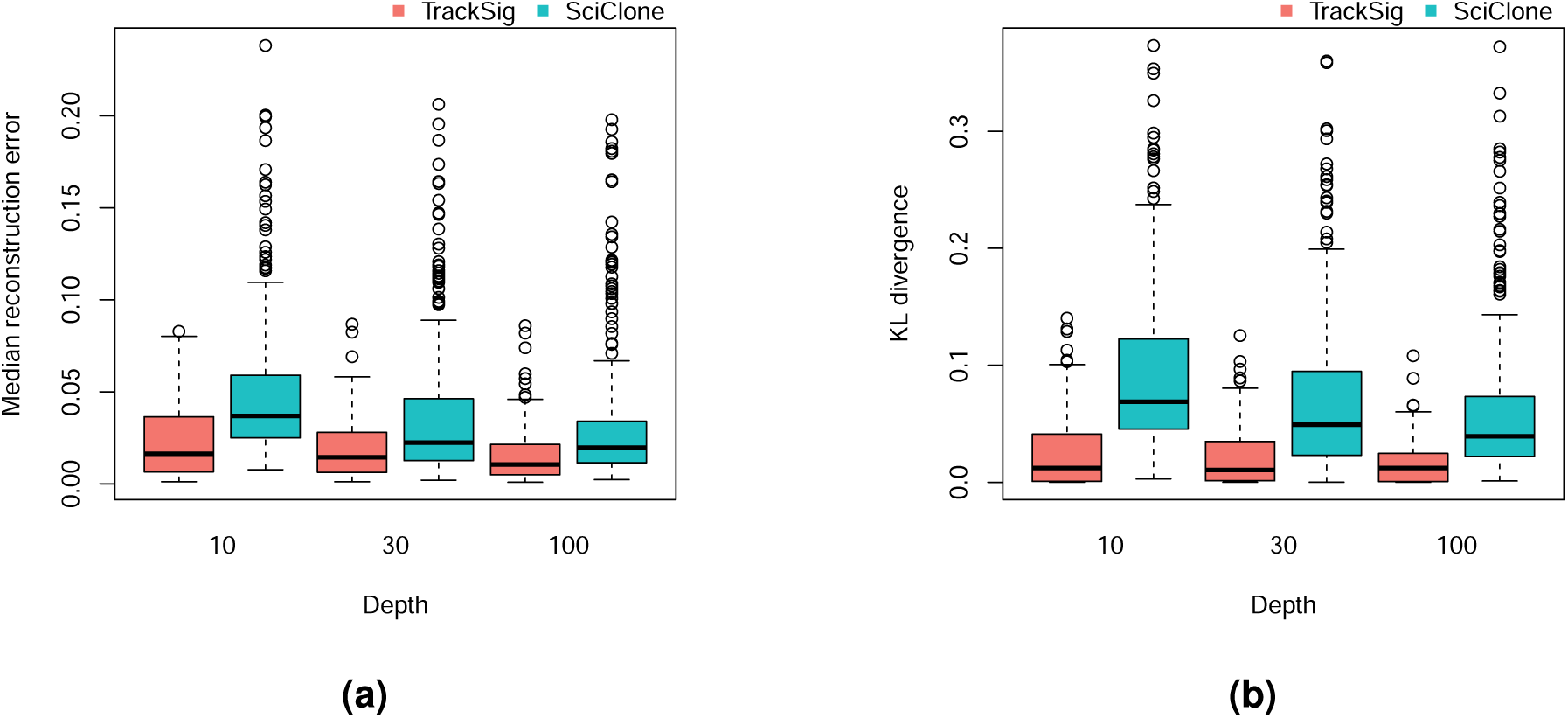
Comparison of TrackSig and SciClone (BMM noise model, default) on one and two cluster simulations with the inclusion of neutral mutations, as described in Appendix C.5. Performance was evaluated across different simulated read depths, shown in X-axis. **(a)** Median absolute, per-mutation difference between true activities and activities estimated by each method. **(b)** Mean, per-mutation, KL divergence between true activities and activites estimated by each method.

**Figure F.9.**
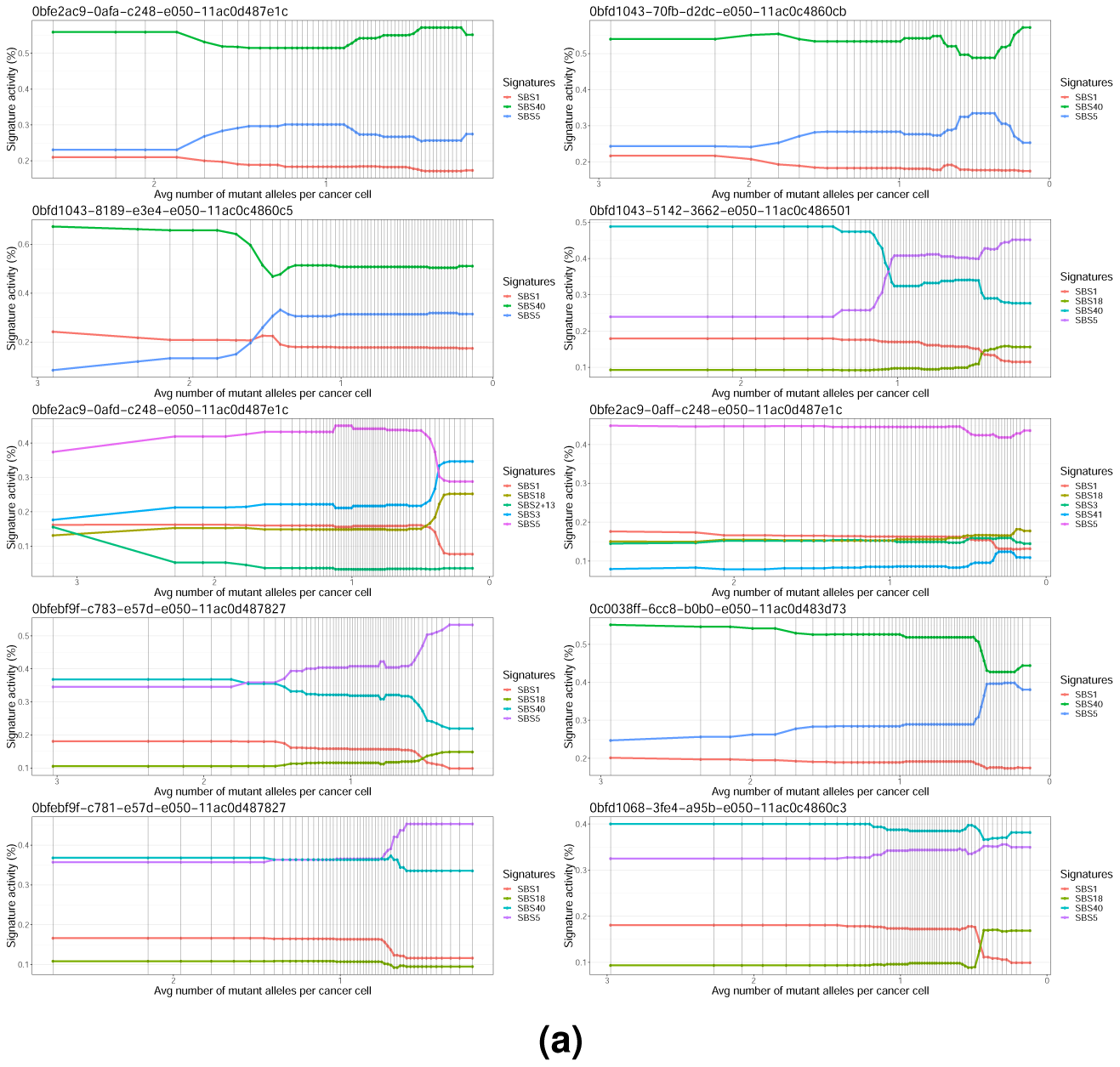
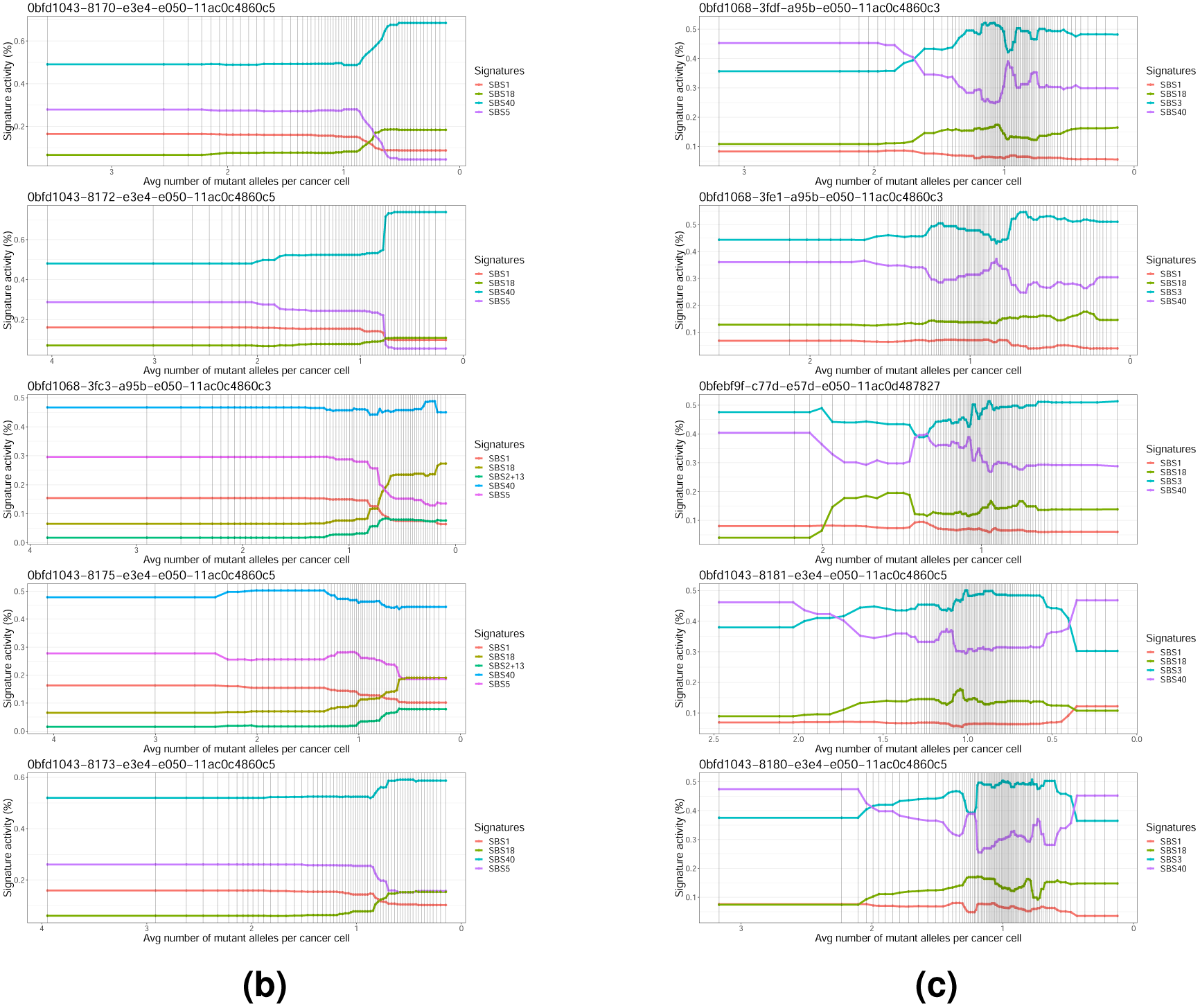
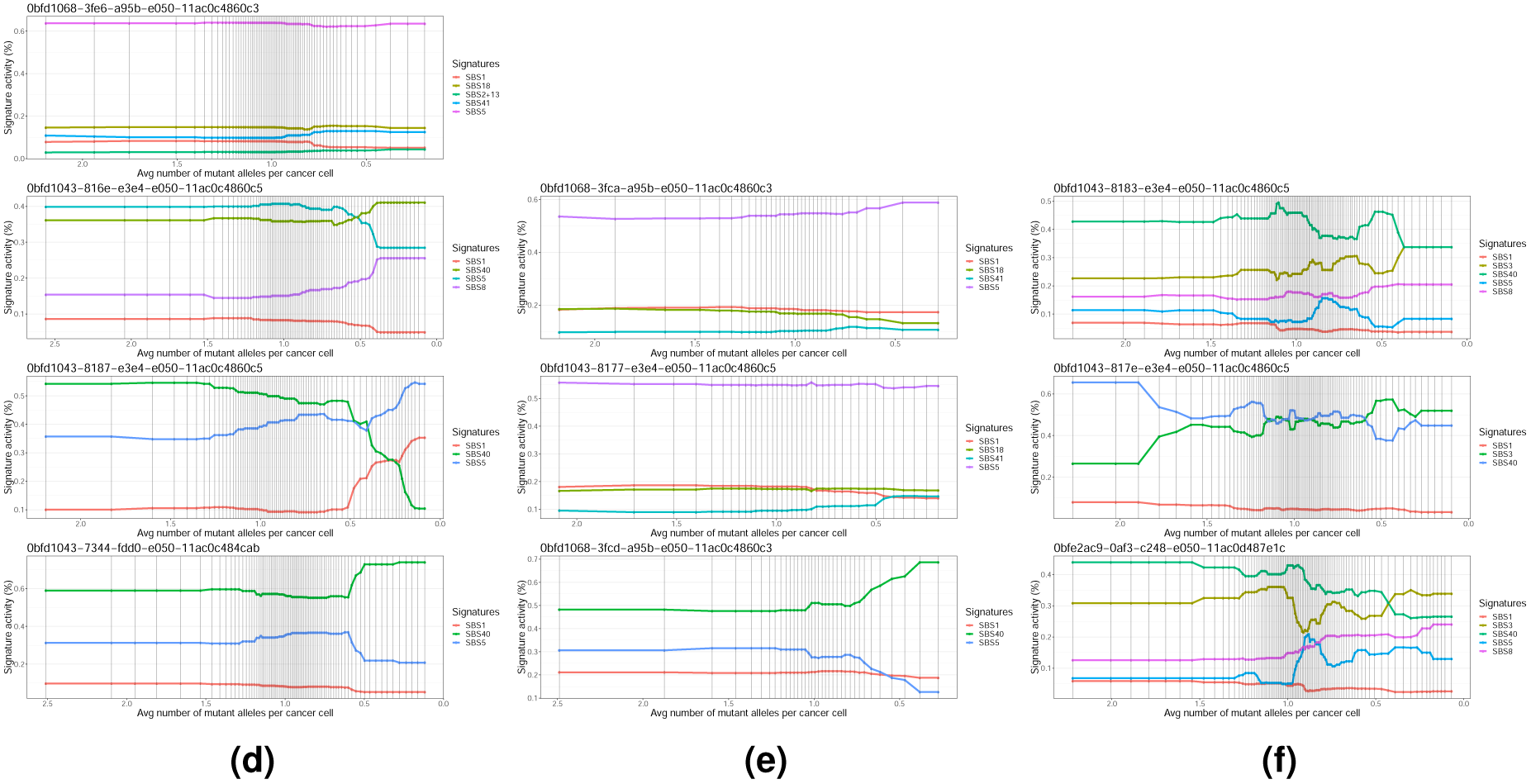
Multi-region cases. Each subplot shows signature trajectories for different samples from the same tumour. Signature trajectories shown are the mean of 30 bootstrap trajectories and therefore are not piece-wise constant. We report mean activity difference and KL divergence between the activities in the *clonal* cluster only. We compare clonal activities across the groups of samples with the same set of active signatures.

**Tumour DO51954:** Group 1 with active signatures “SBS1 SBS5 SBS40”: mean activity diff 0.0573, KL divergence 0.05. Group 2 with signatures “SBS1 SBS5 SBS18 SBS40”: mean activity diff 0.036, KL divergence 0.018

**Tumour DO51958:** Group 1 with active signatures “SBS1 SBS5 SBS18 SBS40”: mean activity diff 0.014, KL divergence 0.002. Group 2 with signatures “SBS1 SBS5 SBS18 SBS40 SBS2+13”: mean activity diff 0.008, KL divergence 0.001.

**Tumour DO51965:** Group with active signatures “SBS1 SBS3 SBS18 SBS40”: mean activity diff 0.042, KL divergence 0.028.

**Tumour DO51953:** Group with active signatures “SBS1 SBS5 SBS40”: mean activity diff 0.031, KL divergence 0.005.

**Tumour DO51959:** Group with active signatures “SBS1 SBS5 SBS18 SBS41”: mean activity diff 0.011, KL divergence 0.0014.

**Tumour DO51962:** Group with signatures “SBS1 SBS3 SBS5 SBS8 SBS40”: mean activity diff 0.037, KL divergence 0.031.

## Footnotes

1 http://cancer.sanger.ac.uk/cosmic/signatures

